# Quantification of the escape from X chromosome inactivation with the million cell-scale human single-cell omics datasets reveals heterogeneity of escape across cell types and tissues

**DOI:** 10.1101/2023.10.14.561800

**Authors:** Yoshihiko Tomofuji, Ryuya Edahiro, Yuya Shirai, Kian Hong Kock, Kyuto Sonehara, Qingbo S. Wang, Shinichi Namba, Jonathan Moody, Yoshinari Ando, Akari Suzuki, Tomohiro Yata, Kotaro Ogawa, Ho Namkoong, Quy Xiao Xuan Lin, Eliora Violain Buyamin, Le Min Tan, Radhika Sonthalia, Kyung Yeon Han, Hiromu Tanaka, Ho Lee, Asian Immune Diversity Atlas Network, Japan COVID-19 Task Force, The BioBank Japan Project, Tatsusada Okuno, Boxiang Liu, Koichi Matsuda, Koichi Fukunaga, Hideki Mochizuki, Woong-Yang Park, Kazuhiko Yamamoto, Chung-Chau Hon, Jay W. Shin, Shyam Prabhakar, Atsushi Kumanogoh, Yukinori Okada

**Affiliations:** Department of Statistical Genetics, Osaka University Graduate School of Medicine, Suita 565-0871, Japan; Integrated Frontier Research for Medical Science Division, Institute for Open and Transdisciplinary Research Initiatives, Osaka University, Suita 565-0871, Japan; Laboratory for Systems Genetics, RIKEN Center for Integrative Medical Sciences, Yokohama 230-0045, Japan; Department of Genome Informatics, Graduate School of Medicine, the University of Tokyo, Tokyo 113-8654, Japan; Department of Respiratory Medicine and Clinical Immunology, Osaka University Graduate School of Medicine, Suita 565-0871, Japan; Genome Institute of Singapore (GIS), Agency for Science, Technology and Research (A*STAR), Singapore 138672, Republic of Singapore; Laboratory for Genome Information Analysis, RIKEN Center for Integrative Medical Sciences, Yokohama 230-0045, Japan; RIKEN Center for Integrative Medical Sciences, Yokohama 230-0045, Japan; Laboratory for Autoimmune Diseases, RIKEN Center for Integrative Medical Sciences, Yokohama 230-0045, Japan; Department of Neurology, Osaka University Graduate School of Medicine, Suita 565-0871, Japan; Department of Infectious Diseases, Keio University School of Medicine, Shinanomachi 160-8582, Japan; Samsung Genome Institute, Samsung Medical Center, Seoul 06351, Korea; Division of Pulmonary Medicine, Department of Medicine, Keio University School of Medicine, Shinanomachi 160-8582, Japan; Department of Pharmacy, National University of Singapore, Singapore 117549, Republic of Singapore; Department of Computational Biology and Medical Sciences, Graduate school of Frontier Sciences, The University of Tokyo, Shirokanedai 108-8639, Japan; Laboratory for Advanced Genomics Circuit, RIKEN Center for Integrative Medical Sciences, Yokohama 230-0045, Japan; Lee Kong Chian School of Medicine, Singapore 308232, Republic of Singapore; Cancer Science Institute of Singapore, Singapore 117599, Republic of Singapore; Department of Immunopathology, Immunology Frontier Research Center, Osaka University, Suita 565-0871, Japan; Laboratory of Statistical Immunology, Immunology Frontier Research Center (WPI-IFReC), Osaka University, Suita 565-0871, Japan; Premium Research Institute for Human Metaverse Medicine (WPI-PRIMe), Osaka University, Suita 565-0871, Japan

**Author notes:** Corresponding authors: Yoshihiko Tomofuji, MD, PhD, Address: Department of Statistical Genetics, Osaka University Graduate School of Medicine, 2-2 Yamadaoka, Suita, Osaka 565-0871, Japan.; Yukinori Okada, MD, PhD, Department of Genome Informatics, Graduate School of Medicine, the University of Tokyo 7-3-1 Hongo, Bunkyo-ku, Tokyo 113-0033, Japan.

## Abstract

One of the two X chromosomes of females is silenced through X chromosome inactivation (XCI) to compensate for the difference in the dosage between sexes. Among the X-linked genes, several genes escape from XCI, which could contribute to the differential gene expression between the sexes. However, the differences in the escape across cell types and tissues are still poorly characterized because no methods could directly evaluate the escape under a physiological condition at the cell-cluster resolution with versatile technology. Here, we developed a method, **<u>s</u>**ingle-**<u>c</u>**ell **<u>L</u>**evel **<u>ina</u>**ctivated **<u>X</u>** chromosome mapping (**scLinaX**), which directly quantifies relative gene expression from the inactivated X chromosome with droplet-based single-cell RNA-sequencing (scRNA-seq) data. The scLinaX and differentially expressed genes analyses with the scRNA-seq datasets of ∼1,000,000 blood cells consistently identified the relatively strong degree of escape in lymphocytes compared to myeloid cells. An extension of **<u>scLinaX</u>** for **<u>multi</u>**-modal datasets, **scLinaX-multi**, suggested a stronger degree of escape in lymphocytes than myeloid cells at the chromatin-accessibility level with a 10X multiome dataset. The scLinaX analysis with the human multiple-organ scRNA-seq datasets also identified the relatively strong degree of escape from XCI in lymphoid tissues and lymphocytes. Finally, effect size comparisons of genome-wide association studies between sexes identified the larger effect sizes of the *PRKX* gene locus-lymphocyte counts association in females than males. This could suggest evidence of the underlying impact of escape on the genotype–phenotype association in humans. Overall, scLinaX and the quantified catalog of escape identified the heterogeneity of escape across cell types and tissues and would contribute to expanding the current understanding of the XCI, escape, and sex differences in gene regulation.

## Introduction

One of the two X chromosomes of females is epigenetically silenced through X chromosome inactivation (XCI) to compensate for the difference in the dosage between sexes. XCI is established on the randomly determined X chromosome in each cell during early embryonic development^1^. Multiple biological processes are involved in the XCI such as the upregulation of the non-coding RNA *XIST*, changes in the histone modifications, and DNA methylation^2^. However, several X-linked genes (∼23% of the X-linked genes^3^) escape from XCI, namely expressed from both active (Xa) and inactive (Xi) X chromosomes.

Expression from Xi due to the escape can contribute to the sex differences of the gene expression and diseases, such as cancer^4^ and autoimmune diseases^5–7^. Furthermore, escape can introduce changes in the effective allele dosage of females in the context of genotype-phenotype association analyses^8–10^ (e.g. genome-wide association study [GWAS] and expression quantitative trait locus [eQTL] mapping). This effect has contributed to the technical difficulties of the X chromosome analyses, resulting in the exclusion of the X chromosome from GWAS and eQTL analyses, which is one of the current limitations of genetic studies. Therefore, understanding XCI escape is important for elucidating the biological sex differences and solving the current limitation of the genetic analysis^11^.

Whether an X-linked gene escapes XCI has historically been determined by evaluating the heterogeneity of metabolic capacity of female cell lines harboring loss of function mutation of X-linked metabolic enzymes on one allele^12,13^. Subsequently, the escape was evaluated for hundreds of genes by analyses of female-derived cell lines with skewed XCI^14^ (i.e. preferential inactivation of the specific X chromosome) and hybridomas from the human and mouse cells^15^. However, concerns remained regarding the generalizability of the findings to physiological conditions within the human body. Although several methods had utilized incomplete XCI skew of the tissue samples for evaluating escape^16–18^, they were often not sensitive and only compatible with samples showing XCI skew.

Differentially expressed gene (DEG) analysis between sexes was also utilized to investigate the escape. For example, DEG analysis of Genotype-Tissue Expression (GTEx) project datasets enabled a comprehensive exploration of the escape in a tissue/gene-wide manner^3^. Although DEG analysis could identify the escape in a physiological condition, it did not directly evaluate the escape and was difficult to separately evaluate the effects of the escape and other factors such as sex-hormonal influences. In addition, previous studies had utilized bulk RNA-seq datasets, thus heterogeneity of the escape across cell types had not been evaluated.

Recently, the single-cell RNA-seq (scRNA-seq) technology has been utilized to analyze the escape from XCI through inference of the Xi and *in silico* generation of the nearly completely skewed XCI condition^3,19,20^. Although scRNA-seq analyses enabled direct observation of the escape under physiological conditions, current computational methods require high per-cell read depth and are compatible only with plate-based scRNA-seq data (e.g. smart-seq). Due to the plate-based method’s relatively limited throughput, analyses have often been performed with a restricted number of samples and cells, and the heterogeneity of the escape across different cell types has remained unexplored. Given that the droplet-based approach (e.g. 10X) is the high-throughput and currently most widely used method, the development of a new method compatible with the 10X dataset is necessary to fully utilize the growing number of publicly available datasets and expand the knowledge of the escape across multiple cell types.

Here, we investigated the escape across immune cell types utilizing the ∼1,000,000 cell-scale 10X peripheral blood mononuclear cells (PBMC) scRNA-seq datasets. We performed pseudobulk and single-cell level DEG analysis to evaluate the escape across cell types. To directly and quantitatively evaluate the escape, we developed a new method **<u>s</u>**ingle-**<u>c</u>**ell **<u>L</u>**evel **<u>ina</u>**ctivated **<u>X</u>** chromosome mapping (**scLinaX**), which identified a heterogeneity of the escape across cell types. We also developed an extension for the multiome (RNA + assay for transposase-accessible chromatin [ATAC]) dataset, **scLinaX-multi**, to evaluate the escape at the chromatin accessibility level. Our scLinaX analysis with a multi-organ dataset, Tabula Sapiens^21^, identified the heterogeneity of the escape across tissues and cell types. Finally, utilizing the quantitative estimates of the escape, we evaluated the effect sizes of sex-stratified eQTL and GWAS analysis to understand how the escape would affect the results of the genotype–phenotype association analyses. scLinaX and scLinaX-multi are publicly available as an R package (https://github.com/ytomofuji/scLinaX).

## Results

### Pseudobulk and single-cell level DEG analysis from the scRNA-seq data of PBMC

To investigate the escape in immune cells, we utilized scRNA-seq data of PBMC generated in the Asian Immune Diversity Atlas project (**Figure 1a**, **Supplementary Table 1**, *N* = 498, 896,511 cells; AIDA) which are derived from healthy Asian subjects. We also utilized previously published PBMC scRNA-seq data (**Supplementary Fig. 1a**, **Supplementary Table 1**; *N* = 147, 865,238 cells) derived from COVID-19 patients and healthy subjects of Japanese ancestry^22,23^.

**Figure 1.**
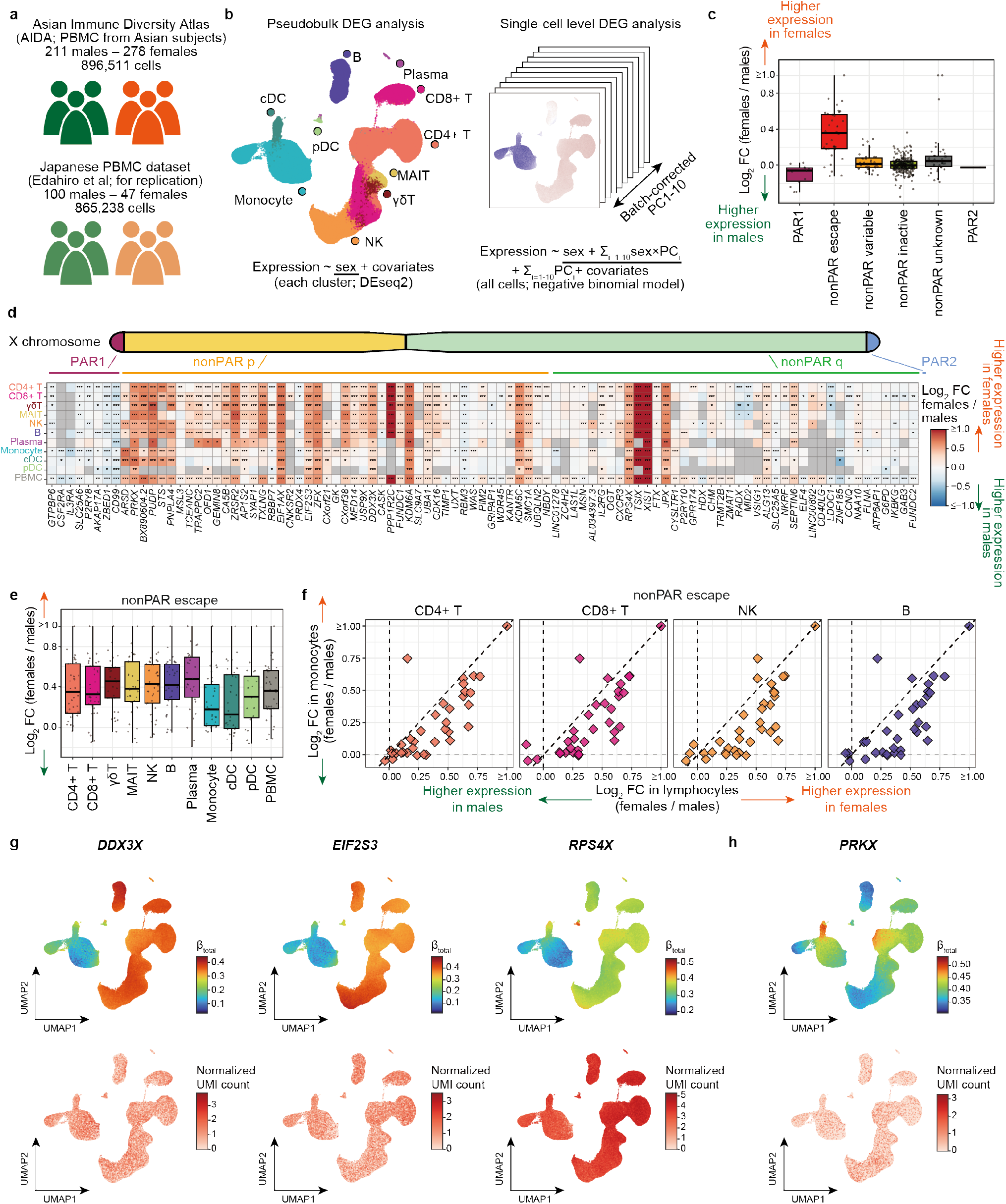
Pseudobulk and single-cell level differentially expressed gene analyses suggested the escape from XCI across immune cells. **a,** Description of the scRNA-seq datasets used in this study. **b,** Description of the DEG analysis methods used in this study. In the pseudobulk DEG analysis, differences in the gene expression level between sexes are evaluated for each cell cluster indicated in the UMAP of the AIDA dataset. In the single-cell level DEG analysis, the difference of the cell states across single cells are represented as continuous batch-corrected PCs and cell-state dependent sex-biased gene expression is evaluated as interactions between the batch-corrected PCs and sex. **c,** A box plot represents log2 fold-changes of the gene expression between sexes. Genes are grouped according to the XCI status annotated in the previous study. The boxplot indicates the median values (center lines) and IQRs (box edges), with the whiskers extending to the most extreme points within the range between (lower quantile − [1.5 × IQR]) and (upper quantile + [1.5 × IQR]). **d,** A heatmap represents differential gene expression between sexes. The colors of the tiles represent log2 fold-changes of the gene expression between sexes. Only genes that satisfied Bonferroni-corrected significance thresholds at least in a cell type are shown. * *P* < 0.05. ** per-cell type FDR < 0.05. *** Bonferroni-corrected *P* < 0.05. **e,** A box plot represents log2 fold-changes of the escapee gene expression between sexes across cell types. The boxplot indicates the median values (center lines) and IQRs (box edges), with the whiskers extending to the most extreme points within the range between (lower quantile − [1.5 × IQR]) and (upper quantile + [1.5 × IQR]). **f,** Scatter plots represent pairwise comparisons of the log2 fold-changes of the escapee gene expression between sexes. The y-axes represent the log2 fold-changes in monocytes and the x-axes represent the log2 fold-changes in lymphocytes. The dashed lines represent *x* = 0, *x* = *y*, and *y* = 0. **g,h,** UMAPs represent the per-cell effect sizes of the sex in the single cell-level DEG analysis calculated as a sum of the effect sizes of sex and sex × batch-corrected PCs (**Methods**, top) and gene expression (bottom). Genes that show a relatively stronger degree of escape in lymphocytes than monocytes (g) and other patterns of heterogeneity of effect sizes (h) are indicated. P-values for the interaction between sex and batch-corrected PCs were < 1 × 10^-200^ (g) and 1.5 × 10^-12^ (h). DEG, differentially expressed genes; FDR, false discovery ratio; IQR, interquartile range; PC, principal component; PAR, pseudoautosomal region; PBMC, peripheral blood mononuclear cells; scRNA-seq, single-cell RNA-seq; UMAP, Uniform manifold approximation and projection; XCI, X chromosome inactivation.

To evaluate the escape across immune cell types, we performed DEG analysis between sexes for each cell type (**Fig. 1b**). Cell types with a large number of cells tended to have a large number of significant DEGs (**Supplementary Fig. 1b**). X-linked genes were enriched among the significant DEGs (*P*_Fisher_ < 0.05 / 11 and *P*_Fisher_ < 0.05 / 8 across cell types, respectively for the two datasets; **Supplementary Fig. 1c**). The results of the DEG analyses were consistent across the two datasets (**Supplementary Fig. 1d,e**). We compared the effect sizes of the X-linked genes in the DEG analysis across the XCI statuses defined in the previous study^3^ and confirmed that known escapee genes tended to have larger effect sizes than other classes of X-linked genes (**Fig. 1c, Supplementary Fig. 1f**). Consistent with the previous study^3^, DEG profile of the X-linked genes is often shared across immune cells (**Fig. 1d**). However, lymphocytes tended to show larger effect sizes than myeloid cells, suggesting the difference of the degree of escape among the immune cells (**Fig. 1e,f, Supplementary Fig. 1g,h**).

To further elucidate the heterogeneity of the female-biased expression of escapee genes among immune cells, we performed single cell-level DEG analysis. We used batch-corrected PCs as proxies for continuous cell state and evaluated the interaction between the sex and cell state using a negative binomial model (**Fig. 1b, Methods**). Significant cell state-interacting sex-biased expression was frequently observed for the escapee genes (**Supplementary Fig. 2a**). The negative binomial model was well-calibrated and the results were consistent across the two datasets (**Supplementary Fig. 2b-d**). The larger effect sizes were observed for the lymphocytes in comparison to the myeloid cells for the representative escapee genes (**Fig. 1g**). On the other hand, some of the escapee genes, such as the *PRKX* gene, showed different patterns of heterogeneity of the effect sizes (**Fig. 1h)**. Overall, heterogeneity of the escape across immune cell types, namely the relatively strong degree of escape in lymphocytes, were suggested from the DEG analysis.

### scLinaX can directly evaluate the escape from the 10X scRNA-seq data

To directly validate the evidence of the heterogeneity of the escape which was indirectly suggested by the DEG analysis, it was effective to directly quantify the escape from XCI, namely gene expression from Xi. 10X scRNA-seq information could be useful for the analysis of escape because single cell-level information enabled us to treat cells with different inactivated X chromosomes separately, while such a method had not been implemented due to the sparse nature of 10X scRNA-seq data. Therefore, we developed a new method, **<u>s</u>**ingle-**<u>c</u>**ell **<u>L</u>**evel **<u>ina</u>**ctivated **<u>X</u>** chromosome mapping (**scLinaX**), which is compatible with the 10X scRNA-seq data (**Fig. 2a**). First, pseudobulk allele-specific expression profiles are generated for cells expressing each candidate reference single nucleotide polymorphism (SNP). Then, alleles of the reference SNPs on the same X chromosome are listed by correlation analysis of the pseudobulk ASE profiles. Finally, scLinaX assigns which X chromosome is inactivated to each cell based on the allelic expression of the reference SNPs and generates a nearly complete XCI skewed condition *in silico* and the estimates for the ratio of the expression from Xi.

**Figure 2.**
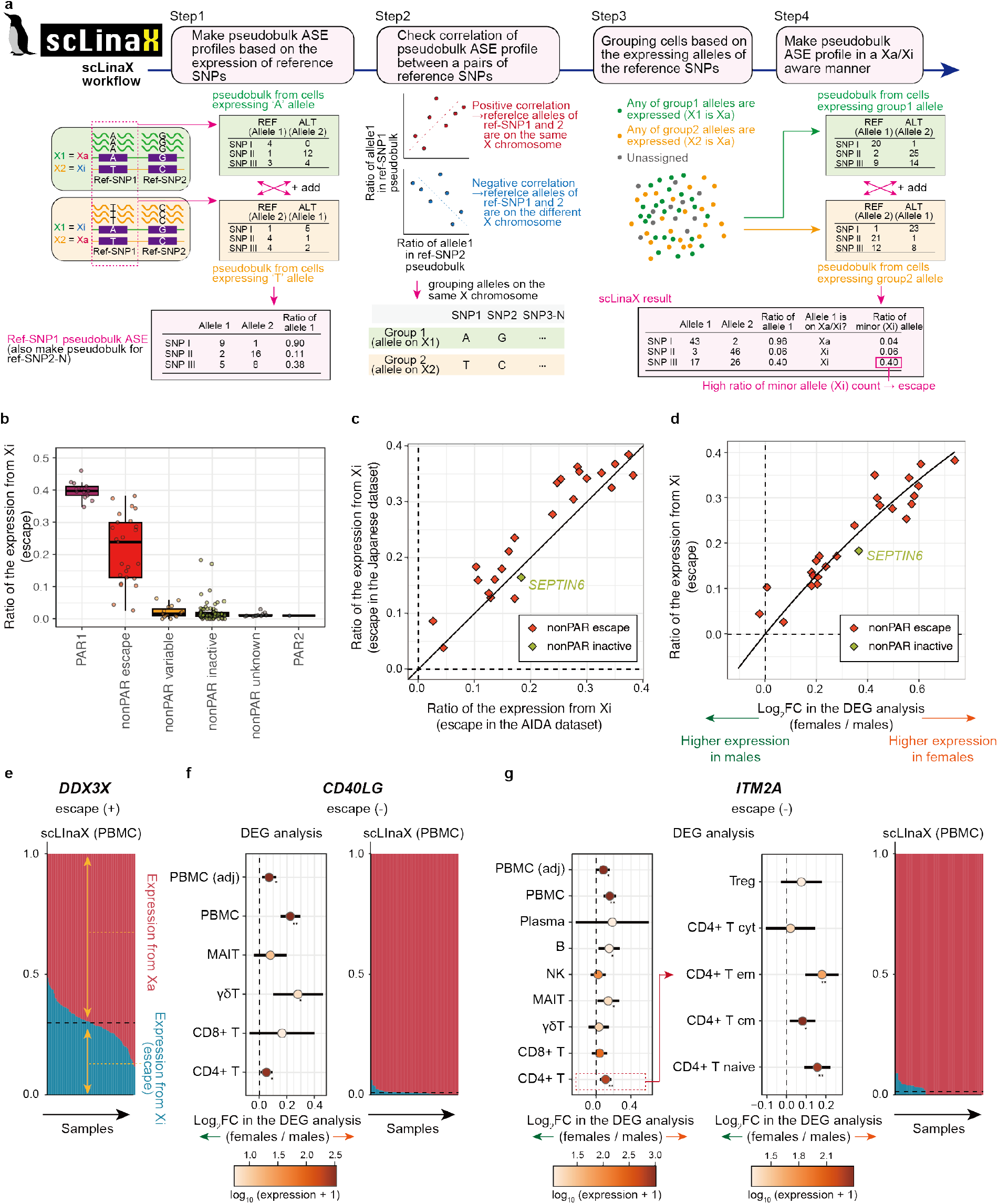
scLinaX, a method to quantify escape from XCI using droplet-based scRNA-seq data. **a,** A schematic illustration of scLinaX. In step1, cells expressing each reference SNP is grouped, and pseudobulk ASE profile are generated. The definitions of alleles 1 and 2 are different across cells depending on which allele of the reference SNP is expressed by each cell. In step2, the correlation between pseudobulk ASE profiles, which are tied to single reference SNPs, is evaluated. Positive and Negative correlation means that the reference alleles of the reference SNPs are on the same strand and different strands, respectively. Based on the results of the correlation analysis, alleles of the reference SNPs are grouped based on which X chromosome are these alleles on. In step3, cells are grouped based on which groups of the reference SNPs are expressed. In step4, pseudobulk ASE profiles from cells expressing any of the reference SNPs are generated. The definition of alleles 1 and 2 are different across cells dependent on which group of the reference SNP allele is expressed by each cell. The ratio of the expression from Xi is defined as a ratio of allele counts from the alleles with a lower allele count. **b,** A box plot represents the estimated ratio of the expression from Xi. Genes are grouped according to the XCI status annotated in the previous study. The boxplot indicates the median values (center lines) and IQRs (box edges), with the whiskers extending to the most extreme points within the range between (lower quantile − [1.5 × IQR]) and (upper quantile + [1.5 × IQR]). **c,** A plot represents the concordance of the ratio of the expression from Xi between the AIDA dataset (x-axis) and Japanese dataset (y-axis). Genes that are annotated as escapee genes and the *SEPTIN6* gene are indicated. The black line indicates *x* = *y*. **d,** A plot represents the relationship between the log2 fold-changes in the DEG analysis (x-axis) and the ratio of the expression from Xi (y-axis). Genes that are annotated as escapee genes and the *SEPTIN6* gene are indicated. The curved line indicates the theoretical relationship under the assumption that differential gene expression between sexes is solely due to the expression from Xi and total gene expression in males and Xa-derived gene expression in females are at the same level. **e,** A plot represents the ratio of the expression from Xa and Xi at an individual level for the *DDX3X* gene. The dashed horizontal line represents the mean ratio of the expression from Xi across samples. **f,g,** Forest plots represent the log2 fold changes in the DEG analysis for each cell type (left) and plots represent the ratio of the expression from Xa and Xi at an individual level (right). The error bars indicate 95% CI. The colors of the dots represent the log-scaled mean normalized count calculated by DEseq2 (baseMean). * *P* < 0.05. ** per-cell type FDR < 0.05. The dashed horizontal line represents the mean ratio of the expression from Xi across samples. AIDA, Asian Immune Diversity Atlas; ALT, alternative allele; ASE, allele-specific expression; CI, confidence interval; DEG, differentially expressed genes; FDR, false discovery ratio; IQR, interquartile range; PAR, pseudoautosomal region; REF, reference allele; SNP, single nucleotide polymorphism; Xa, active X chromosome; XCI, X chromosome inactivation; Xi, inactive X chromosome.

We applied scLinaX to the PBMC single-cell RNA-seq data and SNP array data, and found that previously identified escapee genes tended to show a higher ratio of the expression from Xi than other classes of genes, suggesting that scLinaX had successfully worked (**Fig. 2b, Supplementary Fig. 3a-f, Supplementary Table 2,3)**. We also performed the analysis based on the SNP data called from scRNA-seq data and the results were almost consistent with the results based on the SNP array data (**Supplementary Fig. 3a-i**), suggesting that scLinaX would be also useful when germline genotype data was not available. The scLinaX estimates were consistent between the two datasets, suggesting the robustness of the scLinaX analysis (**Fig. 2c**). Among the genes annotated as subjected to complete XCI, *SEPTIN6* showed a relatively high ratio of the expression from Xi consistently in both of the datasets (**Fig. 2c**, the ratio of the expression from Xi = 0.183 and 0.165 [SD = 0.067 and 0.078], respectively in the AIDA and Japanese datasets). Given that *SEPTIN6* showed female-biased expression in the DEG analysis (log_2_ FC = 0.36 and 0.34 [SE = 0.017 and 0.042], respectively in the AIDA and Japanese datasets) and recently reported to be escapee^18,24^, *SEPTIN6* was thought to actually be an escapee gene.

The relationship between the effect sizes of the DEG analysis and the ratio of the expression from Xi estimated by the scLinaX was compatible with the assumption that differential gene expression between sexes was due to the expression from Xi (**Fig. 2d, Supplementary Fig. 3j**; the ratio of the expression from Xi [y-axis] = 1-1/2^log2^ ^fold^ ^change^ ^[x-axis]^). However, there existed genes that showed female-biased expression in the DEG analysis, but with a low ratio of the expression from Xi. For example, the *CD40LG* gene was female-biased DEG in the PBMC analysis but its ratio of the expression from Xi was low compared to other escapee genes (**Fig. 2e,f**). The *CD40LG* was highly expressed in CD4 T cells, but it was not a DEG in the pseudobulk analysis on CD4 T cells, suggesting that it was detected as a DEG due to confounding of the relative composition of CD4 T cells, not escape (**Fig. 2f, Supplementary Fig. 3k**). The *ITM2A* gene was also detected as a significant female-biased DEG in the PBMC analysis while the ratio of the expression from Xi was low (**Fig. 2g, Supplementary Fig. 3k**). Since *ITM2A* showed significant female-biased expression in the per-cell type DEG analysis, it might be a case that female-biased expression of *ITM2A* was due to the other factors such as sex-hormonal effects. Considering these examples, scLinaX would be useful to directly evaluate the escape and complement the limitation of the DEG analysis.

### Quantification of the escape across cell types by scLinaX

Next, we evaluated the escape as a ratio of the expression from Xi for each cell type by scLinaX. Consistent with the results of the DEG analysis, lymphocytes tended to have a higher ratio of expression from Xi than monocytes for the escapee genes (**Fig. 3a,b, Supplementary Fig. 4a,b, Supplementary Table 2,3**). When per-cell type estimates from scLinaX were projected onto the UMAP, the gradients of the ratio of the expression from Xi showed the same pattern as those from the single-cell level DEG analysis (**Fig. 1g,3c, Supplementary Fig. 4c**). In addition, the *PRKX* gene, which showed an atypical pattern of the heterogeneity of the effect sizes in the DEG analysis, also showed the gradients of the ratio of the expression from Xi with the same pattern as those from the single-cell level DEG analysis (**Fig. 1h,3d, Supplementary Fig. 4c**). Considering the clear relationship between the results of DEG and scLinaX analyses in the bulk PBMC analysis (**Fig. 2d**), these findings suggested that the inter-cell type heterogeneity of the escape quantified by scLinaX contributed to the heterogeneity of sex-difference of the gene expression across cell types.

**Figure 3.**
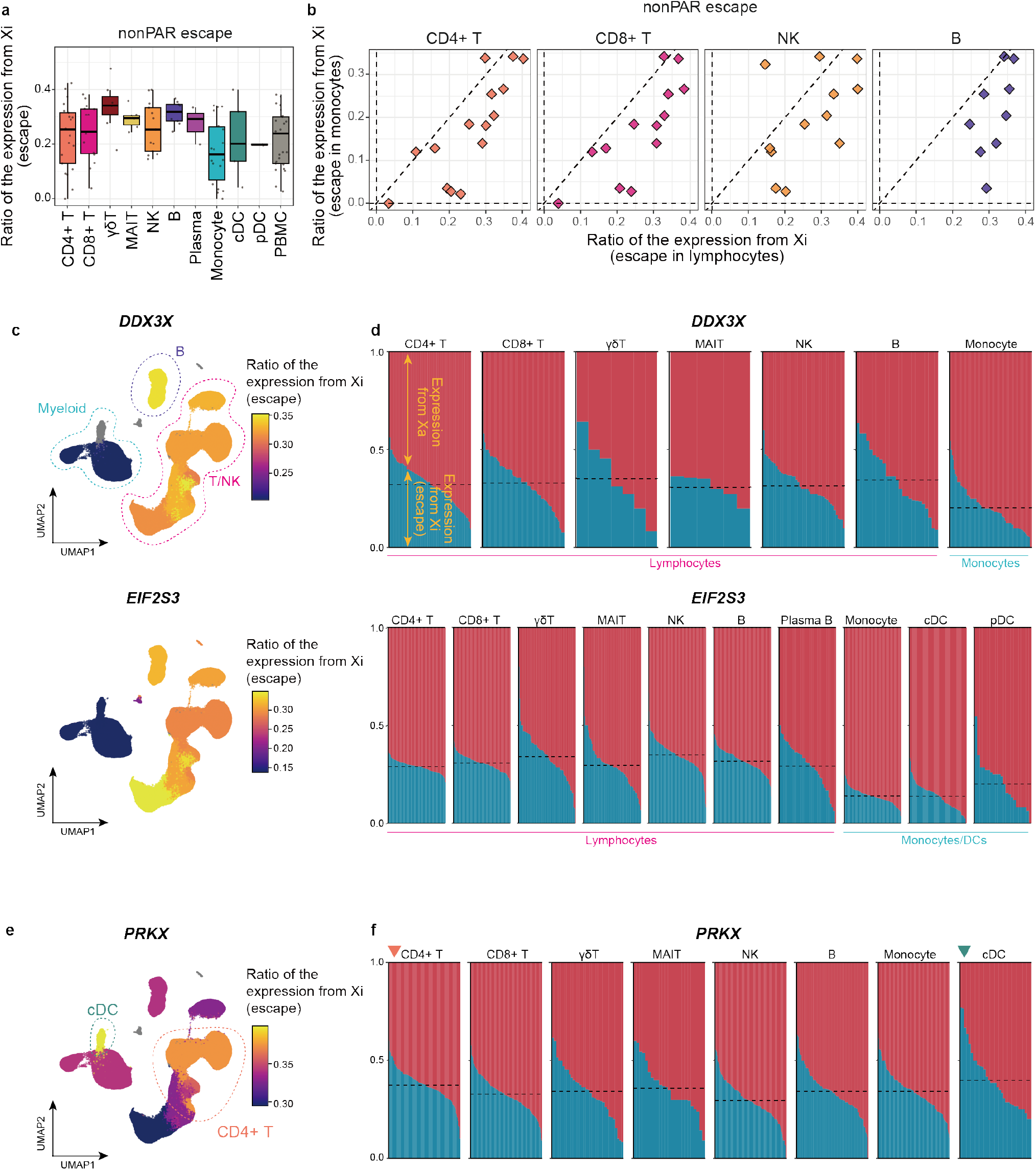
The scLinaX-based quantification of the escape from XCI across immune cell types. **a,** A box plot represents the estimated ratio of the expression from Xi for escapee genes across cell types. The boxplot indicates the median values (center lines) and IQRs (box edges), with the whiskers extending to the most extreme points within the range between (lower quantile − [1.5 × IQR]) and (upper quantile + [1.5 × IQR]). **b,** Scatter plots represent pairwise comparisons of the ratio of the expression from Xi for escapee genes. The y-axes represent the ratio of the expression from Xi in monocytes and the x-axes represent the ratio of the expression from Xi in lymphocytes. The dashed lines represent *x* = 0, *x* = *y*, and *y* = 0. **c,** UMAPs colored according to the ratio of the expression from Xi estimated for each cell type. Representative genes that showed a higher ratio of expression from Xi in lymphocytes than monocytes, the *DDX3X* and *EIF2S3* genes, are indicated. Cell types whose ratio of the expression from Xi could not be estimated are colored grey. **d,** Plot represents the ratio of the expression from Xa and Xi at an individual level for each cell type. Representative genes that show a higher ratio of expression from Xi in lymphocytes than monocytes, the *DDX3X* and *EIF2S3* genes, are indicated. The dashed horizontal line represents the mean ratio of the expression from Xi across samples for each cell type. **e,** A UMAP colored according to the ratio of the expression from Xi estimated for each cell type. The *PRKX* gene, which shows a unique pattern of heterogeneity of the escape across cell types, is indicated. Cell types whose ratio of the expression from Xi could not be estimated are colored grey. **f,** Plot represents the ratio of the expression from Xa and Xi at an individual level for each cell. The *PRKX* gene, which shows a unique pattern of heterogeneity of the escape across cell types, is indicated. The dashed horizontal line represents the mean ratio of the expression from Xi across samples for each cell type. IQR, interquartile range; UMAP, Uniform manifold approximation and projection; Xa, active X chromosome; XCI, X chromosome inactivation; Xi, inactive X chromosome.

### scLinaX-multi can evaluate the escape at the chromatin accessibility level

XCI escape, which we had observed at the transcription level, was closely linked to the gene regulation at the chromatin level. XCI induces chromatin-level transcriptional repression on Xi, while the transcriptionally active chromatin state on Xi can be observed under the escape from XCI. Although previous studies had demonstrated the escape at the chromatin level through the comparative analyses between sexes^25^ and allele-specific epigenetic investigations using cell lines^26^, the chromatin-level escape had not been directly quantified under the physiological condition. To directly quantify the chromatin level escape, we developed an extension of scLinaX for the multi-modal single-cell data (RNA + ATAC), **<u>scLinaX</u>** for **<u>multi</u>**-modal data (**scLinaX-multi**; **Fig. 4a**). In multi-modal single-cell data, each cell has both the RNA and ATAC information. scLinaX-multi utilizes allelic RNA expression information to estimate which X chromosome is inactivated for each cell as done in the scLinaX analysis. For the cells successfully estimated for the inactivated X chromosome based on the RNA information, allelic ATAC information is utilized to calculate the ratio of the accessible chromatin derived from Xi, namely the escape at the chromatin accessibility level.

**Figure 4.**
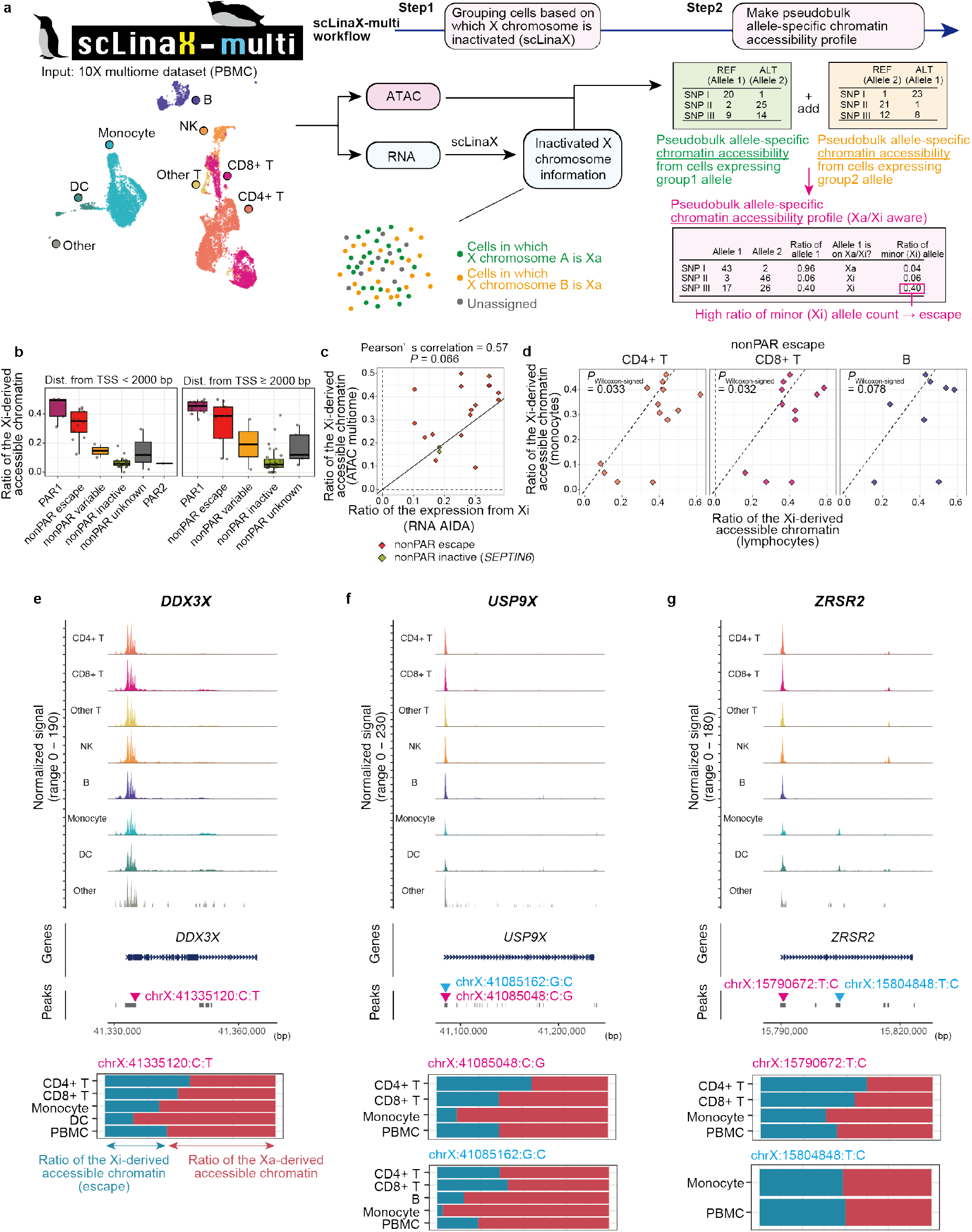
scLinaX-multi, a method to estimate the chromatin accessibility of Xi from multi-modal single-cell omics data. **a,** A schematic illustration of the scLinaX-multi. The input of the scLinaX-multi is single-cell multiome ATAC + Gene Expression data. In step1, cells are grouped based on which X chromosome is inactivated by applying scLinaX to the gene expression information of the 10X multiome data. In step2, pseudobulk allele-specific chromatin accessibility profiles are generated by summing up the allele-specific chromatin accessibility data of each single cell. The definition of alleles 1 and 2 are different across cells dependent on which X chromosome is inactivated in each cell. The ratio of the Xi-derived accessible chromatin is defined as a ratio of allele counts from the alleles with a lower allele count. **b,** Box plots represent the estimated ratio of the accessible chromatin derived from Xi for peaks within 2kbp of TSS (left) and ≥2kbp distant from TSS (right). Peaks are grouped according to the XCI status of the nearest gene. The boxplot indicates the median values (center lines) and IQRs (box edges), with the whiskers extending to the most extreme points within the range between (lower quantile − [1.5 × IQR]) and (upper quantile + [1.5 × IQR]). **c,** A plot represents the relationship between the ratio of the expression from Xi (RNA-level, x-axis) and the ratio of the accessible chromatin derived from Xi (y-axis) for each peak–nearest gene pair. Genes that are annotated as escape genes or showed evidence of escape in the scLinaX analysis (ratio of the expression from Xi > 0.15) are indicated. The black line indicates *x* = *y*. When a single gene has multiple peaks, the average across the peaks for the ratio of the Xi-derived accessible chromatin is used for the calculation of Pearson’s correlation. **d,** Scatter plots represent pairwise comparisons of the accessible chromatin derived from Xi for peaks whose nearest genes are escapee genes. The y-axes represent the ratio of the expression from Xi in monocytes and the x-axes represent the ratio of the expression from Xi in lymphocytes. The dashed lines represent *x* = 0, *x* = *y*, and *y* = 0. P-values are calculated by the Wilcoxon signed-rank test. **e,f,g**, The results of the scLinaX-multi for the representative peaks around escapee genes, namely *DDX3X* (e), *USP9X* (f), and *ZRSR2* (g). Normalized tag counts across cell types are indicated with peak information (top). The ratio of the accessible chromatin derived from Xa and Xi across cell types is indicated as bar plots (bottom) with information on which SNPs are used for the analysis. AIDA, Asian Immune Diversity Atlas; ALT, alternative allele; ATAC, Assay for Transposase-Accessible Chromatin; PAR, pseudoautosomal region; SNP, single nucleotide polymorphism; REF, reference allele; TSS, transcription start site; Xa, active X chromosome; XCI, X chromosome inactivation; Xi, inactive X chromosome.

We applied scLinaX-multi to the publicly available PBMC multiome datasets from a female and found that peaks whose nearest genes were escapee genes tended to show a higher ratio of the accessible chromatin derived from Xi than other classes of peaks, suggesting that scLinaX-multi had successfully worked (**Fig. 4b, Supplementary Fig. 5a-e, Supplementary Table 4**). The ratio of the accessible chromatin derived from Xi (ATAC) and the ratio of the expression from Xi (RNA) for peak–nearest gene pairs were nominally correlated for the escapee genes in PBMC (**Fig. 4c, Supplementary Fig. 5f**; Pearson’s correlation = 0.57 and *P* = 0.066). The ratio of the accessible chromatin derived from Xi was nominally higher in lymphocytes than in monocytes (**Fig. 4d**, *P*_Wilcoxon-signed_ < 0.05 in CD4+ T cells vs. monocytes and CD8+ T cells vs. monocytes). For example, peaks at the transcription start sites (TSS) of the escapee genes (*DDX3X*, *USP9X*, and *ZRSR2*) showed a relatively higher ratio of the accessible chromatin derived from Xi in lymphocytes than in monocytes (**Fig. 4e-g**). In addition, we found the chromatin-level escape at the myeloid cell-specific enhancer in the *ZRSR2* gene locus which were also defined as a cis-regulatory elements (cCRE) in the ENCODE project (EH38E3926410)^27^. In summary, scLinaX-multi could be useful in identifying chromatin-level escape and its heterogeneity across cell types.

### Direct quantification of the escape across multi-organs with scLinaX

To evaluate the heterogeneity of the escape beyond blood cells, we applied scLinaX to the Tabula Sapiens^21^, the current largest publicly available human multi-organ scRNA-seq dataset in terms of number of cells and organs^21^ (https://tabula-sapiens-portal.ds.czbiohub.org). Although the Tabula Sapiens dataset did not contain genotype data, scLinaX was applicable to datasets without genotype data (**Supplementary Fig. 3a-i**). Data from 6 females were included in the analysis, and escapee genes were shared across the organs (**Fig. 5a, Supplementary Fig. 6a-g, Supplementary Table 5**), consistently with the previous study^3^. To evaluate the heterogeneity of the escape across organs, we performed pairwise comparisons of the ratio of the expression from Xi and found that lymphoid tissues, such as lymph node, thymus, and spleen, had a relatively high ratio of the expression from Xi (**Fig. 5b,c**).

**Figure 5.**
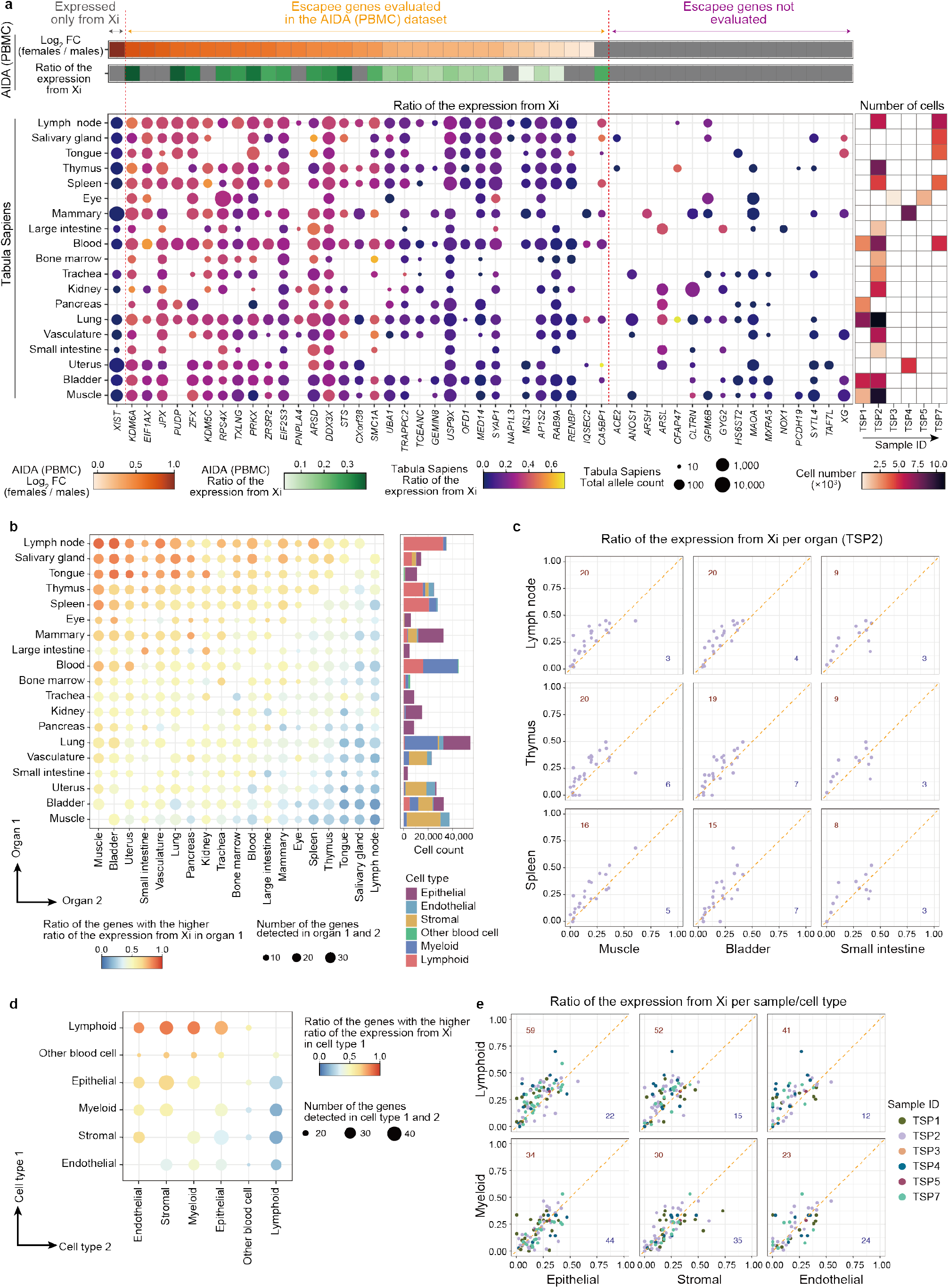
Quantitative evaluation of the escape from XCI with a human multi-organ atlas of single-cell transcriptome. **a,** A dot plot represents the ratio of the expression from Xi across organs from the Tabula Sapiens dataset (y-axis) for escapee genes (x-axis). The color and size of the dots represent the ratio of the expression from Xi and the total allele count. Heatmaps placed above the dot plot represent the log2 fold-change of gene expression between sexes (orange) and the ratio of the expression from Xi (green) calculated from the AIDA dataset. A heatmap placed on the right of the dot plot represents the number of cells used for the scLinaX analysis across organs and samples. For the *XIST* gene, the color of the dots exceptionally represents the expression from Xa. **b,** A dot plot represents the results of the pairwise comparison of the ratio of the expression from Xi across organs. The color of the dots represents the ratio of the genes whose ratio of the expression from Xi is higher in organ 1 (y-axis) than in organ 2 (x-axis). The size of the dots represents the number of genes that are used for each comparison. A bar plot placed on the right of the dot plot represents the numbers and types of the cells which are used for the scLinaX analysis. **c,** Scatter plots represent pairwise comparisons of the ratio of the expression from Xi for escapee genes. The y-axes represent the ratio of the expression from Xi in lymphoid tissues and the x-axes represent the ratio of the expression from Xi in organs with a relatively weak degree of escape. Since these organs are commonly evaluated in TSP2, data from TSP2 is presented. The dashed line represents *x* = *y*. The described numbers indicate the number of genes that are located in the x < y (lower right, blue) and x < y (upper left, red). **d,** A dot plot represents the results of the pairwise comparison of the ratio of the expression from Xi across cell types. The color of the dots represents the ratio of the genes whose ratio of the expression from Xi is higher in cell type 1 (y-axis) than in cell type 2 (x-axis). The size of the dots represents the number of genes that are used for each pairwise comparison. **e,** Scatter plots represent pairwise comparisons of the ratio of the expression from Xi for each escapee gene and individual. The y-axes represent the ratio of the expression from Xi in immune cell types and the x-axes represent the ratio of the expression from Xi in other cell types. The color of the points represents each sample. The dashed line represents *x* = *y*. The described numbers indicate the number of genes–sample pairs that are located in the x < y (lower right, blue) and x < y (upper left, red). AIDA, Asian Immune Diversity Atlas; PBMC, peripheral blood mononuclear cells; Xi, inactive X chromosome; XCI, X chromosome inactivation.

In the analyses of PBMC, it had been suggested that lymphocytes showed relatively strong escape. Therefore, we hypothesized that the relatively high ratio of the expression from Xi observed in the lymphoid tissues was due to the high cell composition of the lymphocytes. Consistent with the hypothesis, a relatively higher ratio of the expression from Xi was observed for the lymphocytes in the pairwise comparisons of the ratio of the expression from Xi across cell types in the Tabula Sapiens dataset (**Fig. 5d,e, Supplementary Table 6**). In summary, scLinaX analysis suggested a tissue-level escape heterogeneity linked to cell type-level escape heterogeneity.

### Evaluation of the differential escape in disease conditions

It was reported that some of the autoimmune diseases (e.g. systemic lupus erythematosus [SLE])-associated genes were escapee and the escape of such genes could be enhanced in the patients with SLE^5–7,28^. Despite the potential association between the escape and diseases, X chromosome-wide evaluation of the escape in disease conditions had not been performed. We analyzed the changes in escape in two diseases, COVID-19^22^ and SLE^29^, based on the scLinaX estimates. After multiple-test correction, we could not detect a significant association possibly because of the lack of power, suggesting the need for future larger cohort analyses (**Supplementary Fig. 7a,b, Supplementary Table 7**). The top nominal association was the increase in the escape of the *EIF2S3* gene in the monocytes of the COVID-19 patients (**Supplementary Fig. 7c**). In COVID-19 patients, *EIF2S3* in monocytes tended to be down-regulated (**Supplementary Fig. 7d**). Therefore, the potential increase of the escape may compensate for the decrease in *EIF2S3* caused by the disease (**Supplementary Fig. 7d**). We also evaluated the escape in a male sample which showed a karyotype of XXY, and the escape status was almost consistent with the healthy females (**Supplementary Fig. 7e, Supplementary Table 8**).

### Difference in the genetic effects on the complex traits was observed at the escapee gene loci

Although genetic association studies such as GWAS and eQTL mapping have successfully identified the genetic backgrounds of human traits, the sex-associated difference is one of the remaining questions. Especially, the X chromosome has been often excluded from the analyses due to technical difficulties despite its apparent importance in the context of sex-associated differences^11^. One of such difficulties is the potential need to adjust the dosage differences between males and females dependent on the degree of the escape for obtaining the per-allele estimate of the GWAS effect sizes. For example, previous literature suggested that the effective dosage of the alleles should be 0/2 for males and 0/1/2 for females under the complete XCI and 0/1 for males and 0/1/2 for females under the complete escape^9^. On the other hand, a previous study showed that the inter-sex differences in the eQTL effects of escape genes were consistent with the complete XCI rather than escape in most cases^8^. Therefore, we evaluated the effects of the escape on the sex differences of the genotype– phenotype association analyses with the quantified catalog of the escape.

First, to evaluate the effects of the escape on the eQTL analysis, we performed eQTL mapping with all samples from the AIDA dataset (allele dosages of the males and females were 0/2 and 0/1/2, respectively) and found 202 significant eQTL signals across 10 cell types (**Supplementary Table 9**; *P* < 5 × 10^-8^). These eQTL signals were highly reproducible by the analysis with the Japanese dataset (**Supplementary Fig. 8a, Supplementary Table 10)**. Then, we performed eQTL mapping separately for males and females and compared the effect sizes of the significant eQTLs on the X chromosome between sexes. We could not observe apparent female-biased effect sizes across all the XCI statuses including escapees (**Fig. 6a, Supplementary Fig. 8b**). In addition, there was no clear relationship between the sex-associated differences of effect sizes and the degree of escape quantified by the DEG and scLinaX analyses (**Fig. 6b, Supplementary Fig. 8c**). These results were consistent to the previous eQTL study^8^ while contradicting to the other studies utilizing ASE or DEG analyses^3,14^ and results of the DEG and scLinaX analyses in this study. We speculate that the sex differences in effective allele dosage caused by escape do not make sex differences in the eQTL effect because of the transformation of the expression data, such as log-transformation which stabilizes variance and resolves heteroskedasticity (**Supplementary Fig. 8d**).

**Figure 6.**
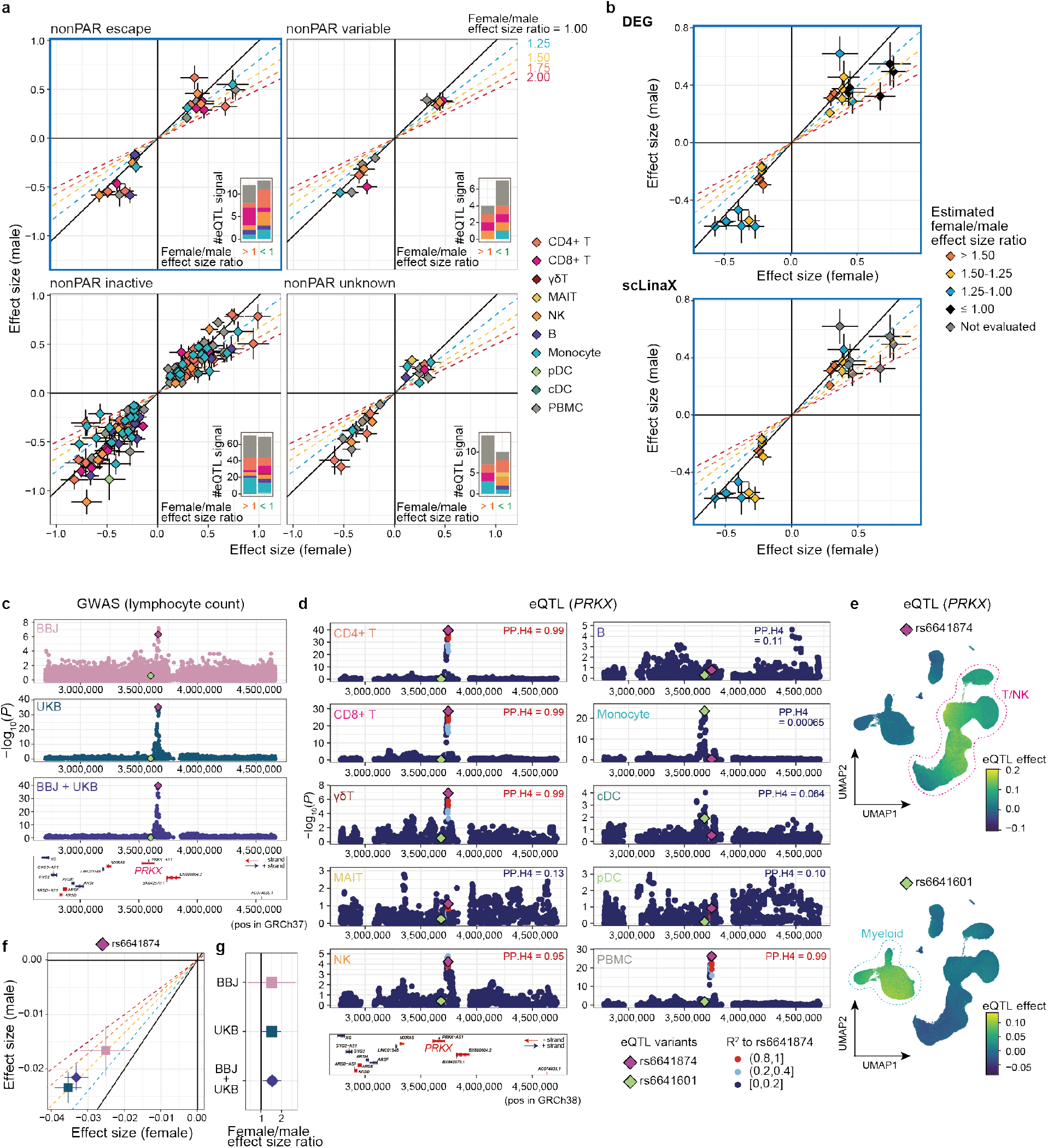
Detection of differential effect sizes between sexes in the genotype– phenotype association analysis. **a,** Scatter plots represent the effect sizes of the significant eQTL signals (*P* < 5 × 10^-8^) in the female-only (x-axis) and male-only (y-axis) analyses, separately for each XCI status. The error bars indicate standard errors. The color of the plots indicates the cell type in which the eQTL signals are identified. The oblique lines correspond to the female/male effect size ratios described in the plots. The attached bar plots indicate the number of eQTL signals that have larger effect sizes in females (left) and males (right). **b,** The scatter plots for escapee genes (a, upper left) are colored according to the estimated female/male effect size ratio based on the DEG analysis (top) and scLinaX analysis (bottom). Genes that are not evaluated in the scLinaX analyses are colored grey. **c,** Locus plots for the association between *PRKX* gene loci and lymphocyte counts in BBJ analysis, UKB analysis, and BBJ + UKB meta-analysis. The x-axis indicates position on the X chromosome (GRCh37) and the y-axis indicates −log10(*P*). The rs6641874 (top variant in the BBJ + UKB meta-analysis and T cells eQTL analysis) and rs6641601 (top variant in the monocytes eQTL analysis) are colored purple and green, respectively. Genes located around the *PRKX* gene region are indicated at the bottom of the plots. **d,** Locus plots for the eQTL analysis of the *PRKX* gene across cell types. The rs6641874 (top variant in the BBJ + UKB meta-analysis and T cells eQTL analysis) and rs6641601 (top variant in the monocytes eQTL analysis) are colored purple and green, respectively. R^2^, a measure for LD to the rs6641874, is indicated as a color of the dots. Results of the colocalization analyses (PP.H4) with lymphocyte counts GWAS in BBJ are indicated in the upper right of the plots. Genes located around the *PRKX* gene region are indicated at the bottom of the plots. **e,** UMAPs represent the per-cell eQTL effect sizes of the variants in the single cell-level eQTL analysis calculated as a sum of the effect sizes of variants and variants × batch-corrected PCs (**Methods**). Associations for *PRKX* genes– rs6641874 (top) and –rs6641601 (bottom) are indicated. P-values for the interaction between genotypes and batch-corrected PCs were 2.7 × 10^-91^ (top) and 2.4 × 10^-51^ (bottom). **f,** Scatter plots represent the effect sizes of the rs6641874 in the female-only (x-axis) and male-only (y-axis) lymphocyte counts GWAS analyses with each cohort. The error bars indicate standard errors. The oblique lines correspond to the female/male effect size ratios described the panel (a). **g,** A forest plot represents the female/male effect size ratios of the rs6641874 in the lymphocyte counts GWAS analyses with each cohort. The error bars indicate 95% CI. BBJ, BioBank Japan; CI, confidence interval; DEG, differentially expressed genes; eQTL, expression quantitative trait locus; GWAS, genome-wide association study; LD, linkage disequilibrium; UKB, UK Biobank; UMAP, Uniform manifold approximation and projection; XCI, X chromosome inactivation.

Next, we evaluated the effects of the escape on the genotype–phenotype association using the two independent biobank datasets. To focus on the genotype–phenotype association signals mediated by the expression of the escapee genes, we evaluated the association between the eQTL variants and blood-related traits using the BioBank Japan (BBJ) dataset (*N* = 82,228-161,145; **Supplementary Table 11,12**)^30,31^. Nine associations satisfied the significance threshold of which only an association between the eQTL variant for *PRKX* (escapee gene) and lymphocyte counts was replicated by the analysis with the UK Biobank (UKB) dataset (**Fig. 6c, Supplementary Fig. 9a,b, Supplementary Table 12**; http://www.nealelab.is/uk-biobank/). Pseudobulk and single-cell level eQTL analyses revealed that two different eQTL signals existed in this region, namely the T/NK cell-specific one and myeloid cell-specific one, and only the T/NK cell-specific eQTL signal colocalized with the GWAS signal (**Fig. 6d,e**). Both of the eQTL signals did not show the difference in the effect sizes between sexes (**Supplementary Fig. 9c**). Interestingly, this locus was suggested to be associated with the white blood cell counts via *PRKX* expression in a female-biased manner in a previous report for the UKB analysis^8^. Given the results of the per-cell type and single-cell level eQTL analysis, this locus could affect the white blood cell counts via the effects on the lymphocytes. Then, we evaluated the effect sizes of the *PRKX* gene loci– lymphocyte counts association in each sex, and found that effect sizes were significantly larger in females than in males (**Fig. 6f,g, Supplementary Table 13**). Although it was difficult to generalize the finding from a single locus, this result might be a piece of evidence for the effect of escape on the difference in the GWAS effect sizes between sexes.

## Discussion

In this study, we quantitatively evaluated the escape from XCI across multiple cell types with large-scale immune-cell and multi-organ scRNA-seq datasets. The newly implemented method, scLinaX, enabled us to directly evaluate the escape across cell types and both the DEG and scLinaX analyses revealed a stronger degree of escape in lymphocytes than myeloid cells. We also implemented an extension of scLinaX for the multi-modal dataset, scLinaX-multi, and revealed a stronger degree of escape in lymphocytes at the chromatin accessibility level. We applied scLinaX to the multi-organ dataset, Tabula Sapiens, and revealed that lymphatic tissues and lymphocytes showed a stronger degree of escape in comparison to other tissues and cell types. Finally, we presented an example of how the escape might have affected sex differences in genotype-phenotype association through the single-cell eQTL analysis and GWAS with two biobank datasets.

scLinaX is a method that enables direct observation of the escape at the cell-cluster level, and its applicability to 10X data makes it highly versatile. Since 10X scRNA-seq data is sparser than plate-based scRNA-seq methods such as smart-seq, single-cell level ASE profiles generated from 10X data are difficult to handle in the same way as plate-based scRNA-seq data. scLinaX resolves the technical difficulty associated with the sparsity of the data by generating pseudobulk ASE profiles for each SNP on the X chromosome and aggregating alleles on the same X chromosome based on the correlation of the pseudobulk ASE profiles of the SNPs. Since the raw output from scLinaX is single-cell level data, it is possible to evaluate the escape in any user-defined cluster including cell types. This unique feature of scLinaX is useful for evaluating the heterogeneity of the escape across various kinds of cells.

scLinaX can map which X chromosome is inactivated for each cell based on the single cell-level transcriptome data, and this information is also useful for evaluating the escape at levels other than the transcriptome level, as demonstrated by the scLinaX-multi analysis with the 10X multiome dataset (RNA + ATAC). The single cell-level multi-modal RNA + ATAC analysis is a relatively new technology and is still in the process of spreading. Therefore, a future generation of the large-scale dataset will enable us to analyze the escape from XCI at the chromatin-accessibility level for a larger number of genes with a million cell-scale dataset. In addition to RNA + ATAC, single-cell joint measurements of RNA + other modalities, such as histone modifications^32^, are currently being developed. Such technologies can enable us to directly observe the escape at the level of the various X chromosome regulations, which will be useful to elucidate the biological mechanisms of the escape.

We identified a unique feature of the lymphocyte, a relatively strong degree of escape through a series of analyses. In a previous analysis utilizing cell imaging, it was revealed that lymphocytes, especially naive ones, had abnormally dispersed distribution of the XIST RNA and reduced normal heterochromatin histone modifications^6,7^. These results suggested that there can be a unique mode of the regulation of XCI in lymphocyte at the chromosome scale. In addition, a relatively strong degree of escape in lymphocytes may also be related to the sex differences in immune phenotype, which could be linked to the higher prevalence of autoimmune diseases in females^33^ and Klinefelter syndrome patients^34^.

How we should handle the allele dosage for males and females and whether allele dosage should be adjusted in the presence of escape is one of the technical difficulties associated with the X chromosome analysis^9,10^. Currently, many software for GWAS, such as PLINK2^35^, BOLT-LMM^36^, and REGINIE^37^, handle the dosage of alleles assuming the complete XCI in a default setting, while previous literature argued that in the presence of escape, the effective dosage on the female should increase^9,10^. In our comparisons of the eQTL effect sizes between sexes, we found no inter-sex differences in eQTL effects regardless of the quantified estimates of the escape. Hence, it might be a case that the effective dosage between sexes could be explained by the sex term in a linear regression model, suggesting that there might not be a necessity to alter the scale of the genotype term in the eQTL analysis of females (**Supplementary Fig. 8d**).

However, this holds true only for a limited trait, such as gene expression, and does not apply to more complex traits contributed by multiple genes. Indeed, in this study, the *PRKX* gene locus was associated with lymphocyte count likely via its eQTL effect in the lymphocytes, and the effect was larger in females than in males. This difference in the effect sizes between sexes might be linked to the increase in allele dosage and *PRKX* expression in females due to escape. Although the limited number of GWAS signal associated with the escapee gene and complexity of the mode of genotype-phenotype associations made it difficult to generalize how the escape affect the sex-difference of the GWAS signal, it would be important to perform GWAS with care for the inter-sex heterogeneity (e.g. sex-stratified analysis^9^). Although the X chromosome has often been excluded from the largest-scale GWAS meta-analyses due to technical difficulties^38,39^, there is a need to actively conduct GWAS of the X chromosome, share sumstats, and promote secondary use in order to overcome this technical difficulty.

In summary, we developed scLinaX, a new method to directly evaluate the escape at the cell-cluster level. We believe that scLinaX and the quantified catalog of escape identified the heterogeneity of escape across cell types and tissues and would contribute to expanding the current understanding of the XCI, escape, and sex differences in gene regulation.

## Methods

### Generation and pre-processing of the AIDA PBMC scRNA-seq data

The Asian Immune Diversity Atlas dataset (v1) was composed of 503 donors of East Asian (Chinese, *N* = 75; Japanese, *N* = 149; Korean, *N* = 165), Southeast Asian (Malay, *N* = 54), and South Asian (Indian, *N* = 60) self-reported ethnicities from Japan, Singapore, and South Korea, and five commercially available European ancestry control samples (LONZA 4W-270). A detailed description of the dataset was included in the flagship manuscript of the Asian Immune Diversity Atlas Network^40^.

The methods for generation and pre-processing of the AIDA PBMC scRNA-seq dataset (v1) are described in the flagship manuscript of the Asian Immune Diversity Atlas Network^40^. Briefly, single-cell RNA-seq for PBMC was performed with 10X Genomics Chromium Controller and 10X Genomics Single Cell 5’ v2 chemistry. We used the DRAGEN Single-Cell RNA pipeline in the Illumina DRAGEN v3.8.4 software (version 07.021.602.3.8.4-20-g74395e76) for pre-processing and genetic demultiplexing. We performed quality control of our dataset in two stages.

We first performed library-level quality control. We started by filtering out cells for which fewer than 300 genes were detected. We then identified the top 2,000 highly variable features using the variance-stabilizing transformation option in Seurat^41^, scaled the data using all genes, and then performed principal component analysis on these highly variable features. We performed nearest-neighbor analyses based on the resulting principal components, and ran Louvain clustering in Seurat at a resolution of 1.0. We annotated the resulting clusters based on a majority vote of the major cell type annotation labels assigned by RCAv2 software^42^ to cells within each cluster. We used the genetic doublet proportion for a library (proportions of mixed genetic identity + ambiguous identity droplets) to estimate the likely total doublet rate for that library^43^. We used this estimate of total doublets in a library, as well as the RCAv2 reference projection-based annotation of clusters (for estimation of homotypic doublet proportion) as part of our input into DoubletFinder^44^, which we used for identifying heterotypic doublets. We then removed cells that had more than 10 (*HBA1* UMIs + *HBB* UMIs), since these cells could be red blood cells, or cells contaminated with red blood cell RNA transcripts.

Then, we performed cell type-specific quality control on our dataset. We removed doublets detected by the DRAGEN genetic demultiplexing workflow and / or DoubletFinder. We then combined single cells from multiple libraries across countries, performed reference projection of such combinations of cells to a reference panel of immune cell transcriptomes using the RCAv2 software^42^, and performed nearest-neighbor analyses based on the principal components of the reference projection coefficients. We performed Louvain clustering and cluster annotation as done in the per-library quality control step. We performed cell type-specific quality control on all single cells across all libraries by applying number of detected genes (including < 300 for platelets, < 500 for myeloid cells, and < 1,000 for other cell types) and percentage mitochondrial reads (> 12.5% for plasma cells and platelets and > 8% for other cell types) filters.

In this study, we removed samples with (i) mismatches between the scRNA-seq inferred sex and reported sex, (ii) < 500 cells per donor, (iii) European genetic ancestry, or (iv) missing/low-quality genotype data. We also removed platelets from the analysis. Finally, we used 896,511 cells from 489 individuals.

### Generation and pre-processing of the PBMC scRNA-seq data of the Japanese healthy and COVID-19 subjects

The PBMC scRNA-seq data of the Japanese was derived from the previously published study^22^. Briefly, peripheral blood samples were obtained from patients with COVID-19 (*N* = 73) and healthy controls (*N* = 75) at Osaka University Hospital. Almost all cases were patients who were transferred from nearby general hospitals because of severe or potentially severe illness during treatment and already initiated with systemic corticosteroid therapy at other hospitals. Single-cell suspensions were processed through the 10X Genomics Chromium Controller following the protocol outlined in the Chromium Single Cell V(D)J Reagent Kits (v1.1 Chemistry) User Guide. Chromium Next GEM Single Cell 5′ Library & Gel Bead Kit v1.1 (PN-1000167), Chromium Next GEM Chip G Single Cell Kit (PN-1000127), and Single Index Kit T Set A (PN-1000213) were applied during the process. Samples were then sequenced on an Illumina NovaSeq 6000 in a paired-end mode.

Droplet libraries were processed using Cell Ranger 5.0.0 (10X Genomics). Filtered expression matrices generated using Cell Ranger count were used to perform the analysis. Cells that had fewer than the first percentile of UMIs or greater than the 99th percentile of UMIs in each sample were excluded. Cells with <200 genes expressed or >10% of reads from mitochondrial genes or hemoglobin genes were also excluded. Additionally, putative doublets were removed using Scrublet (v0.2.1)^45^ and scds (v1.10.0)^46^ for each sample.

The R package Seurat (v4.1.0)^41^ was used for data scaling, transformation, clustering, and dimensionality reduction. Data were scaled and transformed using the SCTransform() function, and linear regression was performed to remove unwanted variation due to cell quality (% mitochondrial reads). For integration, 3,000 shared highly variable genes (HVGs) were identified using SelectIntegrationFeatures() function. Principal component analysis (PCA) was run on gene expression, followed by batch correction using harmony (v0.1)^47^. UMAP dimension reduction was generated based on the first 30 harmony-adjusted principal components. A nearest-neighbor graph using the first 30 harmony-adjusted principal components was calculated using FindNeighbors() function, followed by clustering using FindClusters() function.

Cellular identity was determined by finding DEGs for each cluster using the FindMarkers() function with parameter ‘test.use=wilcox’, and comparing those markers to known cell type-specific genes. Two rounds of clustering were performed (1st, all cells; 2nd, separately for monocytes/DC, T/NK cells, and B cells) and cell type annotation was assigned at the three layers of the granularity based on the marker gene expression. In this study, we mainly used the coarsest annotation (L1) to maintain the number of cells per cluster. In this study, a male subject with COVID-19 was removed because of the aneuploidy of the X chromosome as done in the original study.

### Generation and pre-processing of the AIDA genotype data

A genotyping of AIDA samples was performed using Infinium Global Screening Array (Illumina). SNPs on the nonPAR X chromosome were treated as diploid in males and heterozygous genotypes of such SNPs were converted into ‘missing’ with PLINK (v1.90b4.4)^48^. Then, we performed quality control of the genotype data with PLINK2 (v2.00a3 9 Apr 2020)^48^. We filtered out samples with a call rate of < 0.98. Note that no samples deviated from the Asian sample clusters in a PCA analysis with the 1,000 Genomes (1KG) Project Phase3v5 samples (*N* = 2,504). We removed variants with a variant call rate of < 0.99, deviation from Hardy–Weinberg equilibrium with *P* < 1.0 × 10^−6^ in each population, or significant allele frequency differences between sexes (*P* < 5.0 × 10^−8^). We also removed the variants whose MAF deviated from the reference panels (|MAF in the AIDA Japanese/Korean/Chinese - MAF in the 1KG EAS | > 0.15, |MAF in the AIDA Indian - MAF in the 1KG SAS | > 0.175, or |MAF in the AIDA Japanese - MAF in the 1KG Japanese | > 0.15). The genotype data after the QC was subjected to the genotype imputation in the Michigan Imputation Server^49^. EAGLE (v2.4)^50^ was used for the haplotype phasing of genotype data and Minimac4 was used for genome-wide genotype imputation. We used the reference panels generated from 1KG Project Phase3v5 samples (N = 2,504) with high coverage (30×) sequencing. We set an imputation quality (R^2^) of 0.3 and 0.7, respectively for the scLinaX analysis and eQTL analysis. We used a relaxed threshold in the scLinaX analysis because the genotype could be also confirmed by the allele information of the scRNA-seq reads. In the eQTL analysis, we removed related samples with PI_HAT > 0.17.

### Generation and pre-processing of the Japanese genotype data

Imputed genotype data for the Japanese dataset was derived from the previously published study^22^. A genotyping of COVID-19 and healthy samples was performed using Infinium Asian Screening Array (Illumina) through collaboration with Japan COVID-19 Task Force (https://www.covid19-taskforce.jp/en/home/). SNPs on the nonPAR X chromosome were treated as diploid in males and heterozygous genotypes of such SNPs were converted into ‘missing’. We applied stringent quality control filters to the samples (sample call rate < 0.98, related samples with PI_HAT > 0.175 or outlier samples from East Asian clusters in PCA with HapMap project samples), and variants (variant call rate < 0.99, deviation from Hardy– Weinberg equilibrium with *P* < 1.0 × 10^−6^, or minor allele count < 5). We also excluded SNPs with > 7.5% allele frequency difference with the representative reference datasets of Japanese ancestry, namely the used the population-specific imputation reference panel of Japanese (*N* = 1,037) combined with 1KG Project Phase3v5 samples (*N* = 2,504)^51,52^ and the allele frequency panel of Tohoku Medical Megabank Project^53^. We used SHAPEIT4 software (v4.2.1)^54^ for the haplotype phasing of genotype data. After phasing, we used Minimac4 software for genome-wide genotype imputation. We used the aforementioned population-specific imputation reference panel of Japanese (*N* = 1,037) combined with 1KG Project Phase3v5 samples (*N* = 2,504). We set an imputation quality (R^2^) of 0.3 and 0.7, respectively for the scLinaX analysis and eQTL analysis. We used a relaxed threshold in the scLinaX analysis because the genotype can be also confirmed by the allele information of the scRNA-seq reads. Since scRNA-seq data was generated in the genome build of GRCh38, we performed a liftover with Picard software.

### Pre-processing of the PBMC 10X multiome data

PBMC 10X multiome data was downloaded from the web repository of the 10X Genomics (https://www.10xgenomics.com/resources/datasets/pbmc-from-a-healthy-donor-granulocytes-removed-through-cell-sorting-10-k-1-standard-2-0-0). The count matrix for the RNA data and fragment data for the ATAC data were jointly processed with the Signac software (v1.9.0)^55^. First, cells satisfying all of the following criteria were kept for the analysis; ATAC tag count < 100,000, ATAC tag count > 25,000, RNA count <25,000, RNA count > 1,000, nucleosome signal < 2, TSS enrichment > 1, percent mitochondrial genes [“^MT-”] < 25, percent hemoglobin genes [“^HB[^(P)]”] < 0.1, and percent platelet genes (PECAM1 and PF4) < 0.25. Then, ATAC peaks were called with macs2 through the CallPeaks() function of the Signac and converted into a count matrix. Putative doublets were removed using DoubletFinder (v2.3.0) and scds (v1.14.0) based on the RNA information. RNA data were scaled and transformed using the SCTransform() function and subjected to a PCA analysis with the top 2,000 highly variable genes. ATAC data was subjected to normalization and dimension reduction based on the latent semantic indexing as implemented in the Signac. Cell type annotation was assigned to each cell by multimodal reference mapping with a Multimodal PBMC reference dataset (https://atlas.fredhutch.org/data/nygc/multimodal/pbmc_multimodal.h5seurat) using the FindTransferAnchors() and TransferData() functions. Cells predicted as platelets or erythrocytes were removed from the analysis. Finally, joint UMAP visualization from RNA (top 50 PCs) and ATAC (top 2-40 LSI components) data was generated by the FindMultimodalNeighbors() function followed by the RunUMAP() function. Peak information was visualized with the CoveragePlot() function in Signac.

### Pre-processing of the scRNA-seq data for a sample with a karyotype of XXY

We used a male sample with a karyotype of XXY who was also in the remission phase of multiple sclerosis. The sample was collected at Osaka University Hospital in the same manner as the Japanese dataset. Library preparation, sequencing, and generation of the count matrix were performed as done for the Japanese dataset. Then a count matrix generated by Cell Ranger 6.0.0 was subjected to a QC with the Seurat R package (v4.3.0). First, cells satisfying all of the following criteria were kept for the analysis; RNA count <25,000, RNA count > 1,000, RNA features > 200, nucleosome percent mitochondrial genes [“^MT-”] < 12, percent hemoglobin genes [“^HB[^(P)]”] < 0.1, and percent platelet genes (PECAM1 and PF4) < 0.25. Putative doublets were removed using DoubletFinder (2.3.0) and scds (v1.14.0) based on the RNA information. RNA data were scaled and transformed using the SCTransform() function and subjected to a PCA analysis with the top 2,000 highly variable genes. Cell type annotation was assigned to each cell by multimodal reference mapping with the Multimodal PBMC reference dataset using the FindTransferAnchors() and TransferData() functions. Cells predicted as platelets or erythrocytes were removed from the analysis.

### Pseudobulk DEG analysis

First, pseudobulk raw UMI count data was generated by aggregating the raw UMI counts from all of the cells for each cell type. Samples with at least five cells were used for the analysis. Then, pseudobulk raw UMI count data was subjected to DESeq2 (v1.38.0)^56^ for the DEG analysis. The formulas for the DEG analysis were the following; gene expression ∼ sex + age + cell count + library (+ cell proportion of the CD4+ T, CD8+ T, gdT, MAIT, NK, B, Plasma B, Monocyte, cDC, and pDC in the cell proportion adjusted analysis; AIDA dataset), gene expression ∼ sex + disease (COVID-19 or healthy control) + age + cell count (Japanese dataset). DEGs were the genes satisfying FDR < 0.05 calculated by the DESeq2. Throughout this paper, annotation from a previous study^3^ was used for the comparative analysis across the XCI statuses.

### Single-cell level DEG analysis

We performed single-cell level regression analysis based on the linear mixed model by modifying the method implemented in a previous study^57^. To represent the continuous state of each cell, we used batch-corrected PCs calculated by harmony (v0.1 for the Japanese dataset) or harmonypy (v 0.0.6 for the AIDA dataset) from the top 30 original PCs. The negative binomial model was fitted with the following formula using glmer.nb() function in the lme4 R library (1.1_31); gene expression (raw UMI count) ∼ sex + age + %mitochondrial gene + log_10_(total UMI count of the cell) + PC1-10 of the raw data + (1 | library) + (1 | individual) (for the evaluation of the main effect with the AIDA dataset), gene expression (raw UMI count) ∼ sex + age + %mitochondrial gene + log_10_(total UMI count of the cell) + PC1-10 of the raw data + batch corrected PC 1-10 + sex × batch corrected PC 1-10 + (1 | library) + (1 | individual) (for the evaluation of the interaction effect with the AIDA dataset), gene expression (raw UMI count) ∼ sex + age + disease + %mitochondrial gene + log_10_(total UMI count of the cell) + PC1-10 of the raw data + (1 | individual) (for the evaluation of the main effect with the Japanese dataset), gene expression (raw UMI count) ∼ sex + age + disease + %mitochondrial gene + log_10_(total UMI count of the cell) + PC1-10 of the raw data + batch corrected PC 1-10 + sex × batch corrected PC 1-10 + (1 | individual) (for the evaluation of the interaction effect with the Japanese dataset). In the evaluation for the main effect, the contribution of the sex to the model was evaluated by the likelihood ratio test. In the evaluation of the interaction effect, the contribution of the sex × batch corrected PC 1-10 to the model was evaluated by the likelihood ratio test. For the calculation of the single-cell level effect sizes of the sex, we summed up the effect sizes of the sex and sex × batch corrected PC 1-10 in the interaction effect analysis as done in the previous study.

### Implementation of scLinaX and scLinaX-multi

#### Generation and QC of the single-cell level ASE profile

First, single-cell level ASE profiles were generated by cellsnp-lite software^58^ (v 1.2.3) for each sample. While cellsnp-lite takes genotype data as input, it can also call genotype data from scRNA-seq data. Therefore, we used imputed genotype data based on the SNP array when available, and used genotype data internally called from scRNA-seq data in other cases. Then, allele frequency and gene information were assigned to the SNPs included in the single-cell level ASE profiles by Annovar (Mon, 8 Jun 2020)^59^, and only the common SNPs (MAF > 0.01 in the matched population of the 1KG dataset; AIDA dataset, EAS and SAS; Japanese dataset, EAS; Tabula Sapiens dataset, ALL; 10X multiome dataset, ALL; Asian sample in the SLE dataset, EAS; European sample in the SLE dataset, EUR; XXY sample, EAS) on the gene (intronic, UTR5, UTR3, exonic, ncRNA_exonic, ncRNA_intronic, and splicing) was retained for the analysis.

#### QC of the candidate reference genes used in scLinaX

In scLinaX, we used SNPs on the genes previously annotated as completely subjected to XCI (nonPAR inactive) as candidates for the reference SNPs^3^. We also set QC criteria for these genes to exclude potentially escaping genes. First, SNPs on nonPAR inactive genes (candidate reference genes) expressed in more than 50 cells were extracted and designated as reference SNP candidates. For each SNP, pseudobulk ASE profiles across all the expressing SNPs were calculated separately for cells expressing the ref allele and alt allele, and these were added together after flipping the ref and alt allele counts for the cells expressing the alt allele. In other words, we made a completely slewed XCI *in silico*. For each sample-reference gene pair, the one with the highest number of cells was retained to remove the redundancy. For the pseudobulk ASE profiles, the SNPs with a total allele count of ≥10 were retained, and the minor allele count ratio was calculated as a ratio of the expression from Xi. The SNPs on the reference gene of each pseudobulk profile were excluded from the pseudobulk profiles to prevent the underestimation of the ratio of the expression from Xi. The following two metrics were then calculated for each candidate reference gene. (1) The average ratio of the expression from Xi for the gene when SNPs on the other candidate reference genes were used as references (2) The average of the ratio of the expression from Xi across the other candidate reference genes when the SNPs on the gene was used as reference. Note that when there were multiple SNPs on the same genes derived from the same sample and reference gene, only one with the highest total allele count was used for the calculation of the metrics. Since there could be a potential escape for genes with high metrics values, we used a threshold of 0.05, 0.075, and 0.1 respectively for the AIDA dataset, Japanese dataset, and SLE dataset, and filtered out the potential escapee genes from the candidate reference SNP list. For the Tabula Sapiens, 10X Multiome, and XXY karyotype data, we used the QC results from the AIDA dataset because there were a relatively small number of samples.

#### Grouping cells based on which X chromosome is inactivated

After defining the candidate reference gene set, we performed the scLinaX analysis. First, SNPs on the candidate reference genes expressed in more than 50 (PBMC scRNA-seq dataset), 30 (10X multiome dataset), or 100 (Tabula Sapiens dataset) cells were extracted for each sample. For each SNP, pseudobulk ASE profiles were calculated separately for cells expressing the ref alleles and alt alleles, and these were added together after flipping the ref and alt allele counts for the cells expressing alt alleles. Then, pseudobulk ASE profiles generated from the same samples were subjected to the pairwise Spearman correlation calculation. We set a threshold for the P-values (< 0.05 for all of the datasets) and correlation coefficients (absolute values > 0.5 for the PBMC datasets and > 0.3 for the Tabula Sapiens dataset) for defining the significant correlations. We generated a group of SNPs that had connected by at least one significant correlation. Then we defined a group of reference SNP alleles on the same X chromosome based on the significant correlations within the group. When assuming the XCI, a significant positive correlation meant that the reference alleles of the two reference SNPs were on the same X chromosomes and a significant negative correlation meant that the reference alleles of the two reference SNPs were on the different X chromosomes. If the contradiction happened during the processing of the correlation information within a group of SNPs (e.g. alternative alleles of the three reference SNPs are predicted to be on the different X chromosomes), such a group of SNPs was removed from the analysis. After defining the group of alleles on the same X chromosome, we divided the cells into three groups; (i) cells expressing only alleles of a group, (ii) cells expressing only alleles of another group, (iii) cells expressing no reference alleles or both groups of the reference alleles.

#### Calculation of the ratio of the expression from Xi

We calculated the pseudobulk ASE profiles across cell groups (i) and (ii) separately and combined them after flipping the ref and alt allele counts for the pseudobulk profiles from group (ii) cells. Then, we calculated the ratio of the expression from Xi as a ratio of the minor allele count under the assumption that the expression from Xi was lower than that from Xa^2^. Only the positions with ≥10 total allele counts were considered. When multiple coding SNPs were detected for a gene in a sample, one with the deepest allele counts was selected to evaluate the ratio of the expression from Xi for the gene. When calculating the ratio of the expression from Xi per cell cluster, pseudobulk ASE profiles were generated from cells within the cell cluster while the definition of the Xi/Xa alleles was based on the pseudobulk ASE profiles from all cells.

#### Summarization of the scLinaX results for the AIDA and Japanese dataset

To obtain the ratio of the expression from Xi for each gene, we calculated the average across the samples that had the coding SNPs with ≥10 total allele counts on that gene. Only the genes for which ≥3 samples were used for calculating the average were considered.

#### Implementation of scLinaX-multi and application to the PBMC 10X multiome data

scLinaX-multi is an extension of scLinaX to the multi-modal dataset. In this study, we estimated which X chromosome was inactivated from the RNA-level information and evaluated the escape at the chromatin accessibility level by using the 10X multiome dataset. First, cells were grouped into the following three groups; (i) cells expressing only alleles of a group, (ii) cells expressing only alleles of another group, (iii) cells expressing no reference SNPs or both groups of the alleles, same as the scLinaX procedure. Then, single-cell level allele-specific chromatin accessibility profiles were generated by cellsnp-lite software. In this study, we used genotype data called from the single-cell ATAC data, while it can also take other types of genotype data. Allele frequency and gene information were assigned to the SNPs included in the single-cell level allele-specific chromatin accessibility profiles and only the common SNPs (MAF > 0.01 in the 1KG ALL dataset) on the ATAC peaks were retained for the analysis. We calculated the pseudobulk allele-specific chromatin accessibility profiles across cell groups (i) and (ii) separately and combined them after flipping the ref and alt allele counts for the pseudobulk profiles from group (ii) cells. Finally, we calculated the ratio of the Xi-derived accessible chromatin as a ratio of the minor allele count. Only the positions with ≥10 total allele counts were considered. When calculating the ratio of the Xi-derived accessible chromatin per cell cluster, pseudobulk allele-specific chromatin accessibility profiles were generated from cells within the cell cluster while the definition of the Xi/Xa allele was based on the pseudobulk allele-specific chromatin accessibility profiles from all cells. When multiple coding SNPs were detected for a peak, one with the deepest allele counts was selected to evaluate the ratio of the Xi-derived accessible chromatin. Exceptionally, when visualizing the escape at the chromatin accessibility level (**Fig. 4f**), we retained both of the SNPs on the peaks at the TSS of the *USP9X* gene.

#### Summarization of the scLinaX results for the Tabula Sapiens dataset

We used the processed Tabula Sapiens dataset contributed by the Tabula Sapiens Consortium (https://tabula-sapiens-portal.ds.czbiohub.org)^21^. For the calculation of the ratio of the expression from Xi, we aggregated the allele counts from Xi and Xa across samples for summarization. The annotation of the organs and cell type was derived from the previous study, while the cell type of ‘immune’ was divided into the ‘Lymphoid’, ‘Myeloid’, and ‘Other blood cell’ considering the difference of the escape across immune cells identified in this study. In the pairwise comparisons of the escape across organs and cell types, genes detected in both organs/cell types 1 and 2 were extracted, and the ratio of the genes with a higher ratio of the expression from Xi in the organ/cell type 1 was used as an indicator of the difference of the escape between the organs/cell types. In addition, comparisons of the ratio of the expression from Xi were performed at the individual level. We used only the TSP2 sample for the evaluation of the difference in the escape across organs because major lymphoid tissues were derived solely from the TSP2.

#### Case-control comparisons of the ratio of the expression from Xi

For the generation of the scRNA-seq bam files of the SLE dataset^29^, we downloaded the fastq files and processed them with Cell Ranger 6.1.2. For the case–control comparisons of the escape from XCI with the COVID-19 and SLE datasets, we considered the coding SNPs with ≥5 total allele counts to increase the sample size. We evaluated the genes (i) considered in ≥5 case samples, (ii) considered in ≥5 control samples, and (iii) the ratio of the expression from Xi calculated from the aggregated allele count data across all samples was ≥0.1. We used a negative binomial model (glm.nb() function in the MASS R library [v7.3_58.1]) to evaluate the case–control differences of the escape using the following formula; allele counts from Xi ∼ disease status + log(total allele count) (offset term).

#### scLinaX analysis with a male sample with a karyotype of XXY

As input genotype data for scLinaX, we used imputed genotype data of the X chromosome (non-PAR region) which were generated and processed in the same manner as the genotype data of the Japanese dataset. Since a single sample was available for this analysis, the ratio of the expression from Xi in the sample was presented as it was.

### Pseudobulk eQTL analysis with the AIDA and Japanese dataset

Raw pseudobulk gene expression data was TMM-normalized and log2-transformed with the edgeR R library (v3.40.0)^60^. The genes with (i) raw UMI count ≥ 5 in more than 20% of the samples and (ii) count per million (CPM) ≥ 0.2 in more than 20% of the samples were filtered out as done in a previous study^61^. Then cis-eQTL was identified by tensorQTL (v1.0.8)^62^ with the ‘--mode cis’ option to obtain the list of the significant eQTL signals and with the ‘--mode cis_nomial’ option to obtain the nominal P-values for all of the gene–cis-variant pairs. tensorQTL was applied for (i) all sample data, (ii) only female data, and (iii) only male data with the ‘--maf_threshold 0.05’ option. Sex (only for all sample data analysis), age, cell count, library, genotype PCs 1-10, and gene expression PCs 1-10 were included as covariates for the AIDA dataset analysis. Sex (only for all sample data analysis), age, disease, cell count, genotype PCs 1-10, and gene expression PCs 1-10 were included as covariates for the Japanese dataset analysis. Genotype PCs were calculated from the SNP array data before imputation by using PLINK2. Gene expression PCs were calculated from the TMM-normalized gene expression data using the prcomp() function in the R. Genotypes of the variants on the X chromosome were coded as 0/1/2 in females and 0/2 in males. We defined eQTL signals satisfying *P* < 5 × 10^-8^ in the AIDA all sample analysis as significant eQTL signals.

### Single-cell level dynamic eQTL analysis

We performed a single-cell level dynamic eQTL analysis based on the linear mixed model by modifying the method implemented in the previous study^57^ to evaluate the heterogeneity of the effects of the eQTL variants (rs6641874 and rs6641601) on the *PRKX* gene expression. As done in the single-cell level DEG analysis, we used batch-corrected PCs calculated by harmonypy from the top 30 original PCs to represent the continuous state of each cell. The negative binomial model was fitted with the following formula using glmer.nb() function in the lme4 R library; gene expression (raw UMI count) ∼ genotype + sex + age + %mitochondrial gene + log_10_(total UMI count) + original PC1-10 of the scRNA-seq data + genotype PC 1-10 + batch corrected PC 1-10 of the scRNA-seq data + genotype × batch corrected PC 1-10 of the scRNA-seq data + (1 | library) + (1 | individual). Genotypes of the variants on the X chromosome were coded as 0/1/2 in females and 0/2 in males. In the evaluation of the interaction effect, the contribution of the genotype × batch corrected PC 1-10 to the model was evaluated by the likelihood ratio test. For the calculation of the single-cell level effect sizes of the eQTL effect, we summed up the effect sizes of the genotype and genotype × batch corrected PC 1-10 of the scRNA-seq data in the interaction effect analysis as done in the previous study.

### GWAS for the blood-related traits with the BBJ cohort

BBJ is a prospective biobank that collaboratively recruited approximately 200,000 patients with ≥1 of 47 diseases and collected DNA, serum samples, and clinical information from 12 medical institutions in Japan between 2003 and 2007. The Japanese samples in BBJ were genotyped with the Illumina HumanOmniExpressExome BeadChip or a combination of the Illumina HumanOmniExpress and HumanExome BeadChips. Quality control of samples and genotypes was conducted as described elsewhere^51^. We analyzed subjects of Japanese ancestry identified by a PCA analysis. Genotype data were imputed with the aforementioned 1KG Project phase3v5 genotype data and Japanese whole-genome sequencing data using Minimac3. As for the blood-related trait data (white blood cell number [WBC], lymphocyte number [LYM], monocyte number [Mono], eosinophils number [EOS], basophils number [BAS], neutrophils number [NEU], hemoglobin [Hb], hematocrit [Ht], mean corpuscular volume [MCV], red blood cell number [RBC], and platelet number [PLT]), we generally used the values measured at the participants’ first visit to the hospitals, and excluded values outside three times the interquartile range (IQR) of the upper or lower quartile across participants as previously described (**Supplementary Table 11**)^31^. Then, blood-related trait data were subjected to the rank-based inverse normal transformation separately for males and females. We conducted X chromosome GWAS for each blood-related trait using REGENIE (v3.2.7)^37^. We included age, sex, and the top 20 principal components as covariates. Genotypes of the variants on the X chromosome were coded as 0/1/2 in females and 0/2 in males.

### Comparisons of the GWAS effect sizes between sexes with the BBJ and UKB cohort

GWAS summary statistics for the UKB cohort were downloaded from the web repository (Nealelab/UK_Biobank_GWAS: v2; Zenodo, https://doi.org/10.5281/zenodo.8011558). Fixed-effect meta-analysis across sexes or cohorts was performed with the metafor R package (v4.2_0). The standard error of the ratio between the female effect sizes (β_female_) and male effect sizes (β_male_) was calculated based on the law of error propagation as previously done^8^.

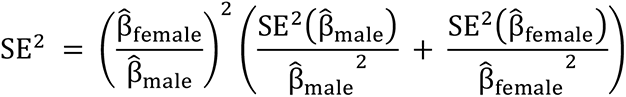

The significance of the difference between the female effect sizes (β_female_) and male effect sizes (β_male_) was evaluated by calculating the following statistics which follow a χ^2^-distribution.

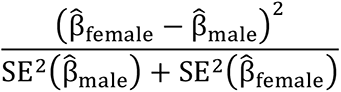

### Evaluation of the colocalization between the GWAS and eQTL signals

To evaluate the colocalization between the lymphocyte count GWAS signals and *PRKX* gene eQTL signals, we used the coloc R package (v5.2.2)^63^. Since the reference human genome was different between the GWAS (GRCh37) and eQTL (GRCh38) analysis, we performed a liftover with the bcftools (v.1.16). Variants within 1,000,000 bp from rs6641874 were used as inputs and PP.H4 > 0.80 was considered as a colocalization of the signals.

## Data availability

The AIDA Data Freeze v1 gene-cell matrix (1,058,909 cells from 503 Japan, Singaporean Chinese, Singaporean Malay, Singaporean Indian, and South Korea Asian donors and 5 distinct Lonza commercial controls), with BCR-seq and TCR-seq metadata, and donor age, sex, and self-reported ethnicity metadata, is available via the Chan Zuckerberg CELLxGENE data portal at https://cellxgene.cziscience.com/collections/ced320a1-29f3-47c1-a735-513c7084d508. The open-access AIDA datasets are available via the Human Cell Atlas Data Coordination Platform at https://data.humancellatlas.org/explore/projects/f0f89c14-7460-4bab-9d42-22228a91f185. Raw scRNA-seq sequencing data for the Japanese dataset are available at the Japanese Genotype-phenotype Archive (JGA) with accession codes JGAS000593/JGAD000722/JGAS000543/JGAD000662^22,23^. All the raw sequencing data of Japanese scRNA-seq dataset can also be accessed through application at the NBDC with the accession code hum0197 (https://humandbs.biosciencedbc.jp/en/hum0197-latest). Genotype data for the Japanese dataset are available at European Genome-Phenome Archive (EGA) with the accession code EGAS00001006950 (https://ega-archive.org/studies/EGAS00001006950).

## Code availability

scLinaX and scLinaX-multi is available as an R package from https://github.com/ytomofuji/scLinaX.

## Acknowledgments

We would like to thank all donors and participants in the studies constituting the Asian Immune Diversity Atlas. The Singapore donor samples were obtained through the Health for Life in Singapore (HELIOS) Study (Lee Kong Chian School of Medicine, Nanyang Technological University; National Healthcare Group, Singapore; Imperial College London). We would like to express our thanks to participants of the HELIOS study and the HELIOS operation team for recruitment, organisation and data/sample collection. This study (NTU IRB: 2016-11-030) is supported by is supported by Singapore Ministry of Health’s (MOH) National Medical Research Council (NMRC) under its OF-LCG funding scheme (MOH-000271-00) and intramural funding from Nanyang Technological University, Lee Kong Chian School of Medicine and the National Healthcare Group. This project has been made possible in part by grant number CZF2019-002446 (to Shyam Prabhakar, Woong-Yang Park, Jay W. Shin, and John Chambers) from the Chan Zuckerberg Foundation, and grant numbers 2020-224570 (to Shyam Prabhakar, Woong-Yang Park, Varodom Charoensawan, Ponpan Matangkasombut, and Partha P. Majumder) and 2021-240178 (to Shyam Prabhakar, Woong-Yang Park, Jay W. Shin, John Chambers, Varodom Charoensawan, Ponpan Matangkasombut, and Partha P. Majumder) from the Chan Zuckerberg Initiative DAF, an advised fund of Silicon Valley Community Foundation. This project was also supported by the Thailand Program Management Unit for National Competitiveness Enhancement (PMU-C) (C10F650132) (to Varodom Charoensawan, Ponpan Matangkasombut, Manop Pithukpakorn, and Bhoom Suktitipat) and Mahidol University’s Basic Research Fund: fiscal year 2021 (BRF1-017/2564) (to Varodom Charoensawan and Bhoom Suktitipat). We would like to thank Jennifer Zamanian, Jennifer Chien, and Jason Hilton from the Human Cell Atlas Lattice team (Stanford University) for their help with and work on data deposits and coordination for community access. B.L. is supported by the Ministry of Education, Singapore, under its Academic Research Fund Tier 1 (FY2023; 23-0434-A0001; 22-5800-A0001) and Tier 2 (MOE-T2EP30123-0015), the Precision Medicine Translational Research Programme Core Funding (NUHSRO/2020/080/MSC/04/PM), NUS ODPRT Seed Funding, and NUS YLLSoM Seed Funding. We want to acknowledge the participants and investigators of BioBank Japan and UK Biobank study. We thank all the members of the Japan COVID-19 Task Force and the Asian Immune Diversity Atlas Network members for their support. We thank Prof. Keishi Fujio at The University of Tokyo and Dr. Mineto Ota at Stanford University for the scientific discussion. We thank Dr. Kazuyoshi Ishigaki and Dr. Masahiro Nakano for kind advises related to the DEG analysis. Y.O. was supported by JSPS KAKENHI (22H00476), and AMED (JP23km0405211, JP23km0405217, JP23ek0109594, JP23ek0410113, JP23kk0305022, JP223fa627002, JP223fa627010, JP233fa627011, JP23zf0127008, JP23tm0524002), JST Moonshot R&D (JPMJMS2021, JPMJMS2024), Takeda Science Foundation, Bioinformatics Initiative of Osaka University Graduate School of Medicine, Institute for Open and Transdisciplinary Research Initiatives, Center for Infectious Disease Education and Research (CiDER), and Center for Advanced Modality and DDS (CAMaD), Osaka University.

## Author contributions

Y.T. and Y.O. designed the study. Y.T., R.E., Y.S., K.K., K.S., Q.S.W., S.N., J.M., Q.L., E.B., R.S., K.H., B.L., and C.C.H. conducted the data analysis. Y.T. and Y.O. wrote the manuscript. R.E., Y.S., and L.T., conducted the experiments. Y.T., R.E., Y.S., K.S., S.N., Y.A., A.S., T.Y., K.O., H.N., H.T., H.L., and T.O. collected and managed the samples. B.L., K.M., K.F., H.M., W.P., K.Y., C.C.H., J.W.S., S.P., A.K., and Y.O. supervised the study. All authors contributed to the article and approved the submitted version.

## Asian Immune Diversity Atlas (AIDA) Network

Atlas assembly authors are arranged by area of contribution and ordered alphabetically by last name.

<u>Single-cell experimental dataset generation leads:</u> Varodom Charoensawan^1,2,3,4,5,6,7^, Chung-Chau Hon^8,9^, Partha P. Majumder^10,11^, Ponpan Matangkasombut^3,12^, Woong-Yang Park^13^, Shyam Prabhakar^14,15,16^, Jay W. Shin^14,17^

<u>Cohort and sample collection leads:</u> Piero Carninci^18,19^, John C. Chambers^15^, Marie Loh^14,15^, Manop Pithukpakorn^6,20^, Bhoom Suktitipat^2,5^, Kazuhiko Yamamoto^21^

<u>Overall study design and protocol development:</u> Deepa Rajagopalan^14^, Nirmala Arul Rayan^14^, Shvetha Sankaran^14^

<u>Sample isolation and processing, single-cell experimental data generation:</u> Juthamard Chantaraamporn^1,2,3^, Ankita Chatterjee^10^, Supratim Ghosh^22^, Kyung Yeon Han^13^, Damita Jevapatarakul^3,12^, Sarintip Nguantad^1,2,3^, Sumanta Sarkar^22^, Narita Thungsatianpun^3,12^

<u>Sample isolation and processing:</u> Mai Abe^21^, Seiko Furukawa^21^, Gyo Inoue^21^, Keiko Myouzen^21^, Jin-Mi Oh^13^, Akari Suzuki^21^

<u>Single-cell experimental data generation:</u> Yoshinari Ando^17,18^, Miki Kojima^18^, Tsukasa Kouno^17^, Jinyeong Lim^13^, Arindam Maitra^22^, Le Min Tan^14^, Prasanna Nori Venkatesh^14^

<u>Single-cell experimental data generation and analysis:</u> Murim Choi^23^, Jong-Eun Park^24^

<u>Single-cell data analysis up to cell type annotation:</u> Eliora Violain Buyamin^14^, Kian Hong Kock^14^, Quy Xiao Xuan Lin^14^, Jonathan Moody^8^, Radhika Sonthalia^14^

<u>Genotype QC and imputation, GWAS summary statistics:</u> Kazuyoshi Ishigaki^25^, Masahiro Nakano^21,26^, Yukinori Okada^27,28,29,30,31,32^, Yoshihiko Tomofuji^27,28,29^

**Affiliations**

1) Department of Biochemistry, Faculty of Science, Mahidol University, Bangkok 10400, Thailand

2) Integrative Computational BioScience (ICBS) Center, Mahidol University, Nakhon Pathom 73170, Thailand

3) Systems Biology of Diseases (SyBiD) Research Unit, Faculty of Science Mahidol University, Bangkok 10400, Thailand

4) Research Department, Faculty of Medicine Siriraj Hospital, Mahidol University, Bangkok 10700, Thailand

5) Department of Biochemistry, Faculty of Medicine Siriraj Hospital, Mahidol University, Bangkok 10700, Thailand

6) Siriraj Genomics, Faculty of Medicine Siriraj Hospital, Mahidol University, Bangkok 10700, Thailand

7) School of Chemistry, Institute of Science, Suranaree University of Technology, Nakhon Ratchasima 30000, Thailand

8) Laboratory for Genome Information Analysis, RIKEN Center for Integrative Medical Sciences, 1-7-22 Suehiro-cho, Tsurumi-ku, Yokohama, Kanagawa 230-0045, Japan

9) Graduate School of Integrated Sciences for Life, Hiroshima University, 1-3-3-2 Kagamiyama, Higashihiroshima, Hiroshima 739-0046, Japan

10) John C. Martin Centre for Liver Research and Innovations, Sonarpur, Kolkata 700150, India

11) Indian Statistical Institute, 203 B.T. Road, Kolkata 700108, India

12) Department of Microbiology, Faculty of Science, Mahidol University, Bangkok 10400, Thailand

13) Samsung Genome Institute, Samsung Medical Center, Seoul 06351, Republic of Korea

14) Genome Institute of Singapore (GIS), Agency for Science, Technology and Research (A*STAR), 60 Biopolis Street, Genome, Singapore 138672, Republic of Singapore

15) Nanyang Technological University, Lee Kong Chian School of Medicine, Clinical Sciences Building, Level 18, 11 Mandalay Road, Singapore 308232, Republic of Singapore

16) Cancer Science Institute of Singapore, National University of Singapore, 14 Medical Drive, Singapore 117599, Republic of Singapore

17) Laboratory for Advanced Genomics Circuit, RIKEN Center for Integrative Medical Sciences, 1-7-22 Suehiro-cho, Tsurumi-ku, Yokohama, Kanagawa 230-0045, Japan

18) Laboratory for Transcriptome Technology, RIKEN Center for Integrative Medical Sciences, 1-7-22 Suehiro-cho, Tsurumi-ku, Yokohama, Kanagawa 230-0045, Japan

19) Genomics Research Center, Fondazione Human Technopole, Viale Rita Levi-Montalcini, 1 -Area MIND, Milano, Lombardy 20157, Italy

20) Department of Medicine, Faculty of Medicine Siriraj Hospital, Mahidol University, Bangkok 10700, Thailand

21) Laboratory for Autoimmune Diseases, RIKEN Center for Integrative Medical Sciences, 1-7-22 Suehiro-cho, Tsurumi-ku, Yokohama, Kanagawa 230-0045, Japan

22) Biotechnology Research and Innovation Council - National Institute of Biomedical Genomics, Kalyani, West Bengal 741251, India

23) Department of Biomedical Sciences, Seoul National University College of Medicine, Seoul 03080, Republic of Korea

24) Graduate School of Medical Science and Engineering, KAIST, Daejeon, Republic of Korea

25) Laboratory for Human Immunogenetics, RIKEN Center for Integrative Medical Sciences, 1-7-22 Suehiro-cho, Tsurumi-ku, Yokohama, Kanagawa 230-0045, Japan

26) Department of Allergy and Rheumatology, Graduate School of Medicine, The University of Tokyo, 7-3-1 Hongo, Bunkyo-ku, Tokyo 113-8654, Japan

27) Laboratory for Systems Genetics, RIKEN Center for Integrative Medical Sciences, 1-7-22 Suehiro-cho, Tsurumi-ku, Yokohama, Kanagawa 230-0045, Japan

28) Department of Statistical Genetics, Graduate School of Medicine, Osaka University, 2-2 Yamadaoka, Suita, Osaka 565-0871, Japan

29) Integrated Frontier Research for Medical Science Division, Institute for Open and Transdisciplinary Research Initiatives, Osaka University, 2-2 Yamadaoka, Suita, Osaka 565-0871, Japan

30) Department of Genome Informatics, Graduate School of Medicine, The University of Tokyo, 7-3-1 Hongo, Bunkyo-ku, Tokyo 113-8654, Japan

31) Laboratory of Statistical Immunology, Immunology Frontier Research Center (WPI-IFReC), Osaka University, 3-1 Yamadaoka, Suita, Osaka 565-0871, Japan

32) Premium Research Institute for Human Metaverse Medicine (WPI-PRIMe), Osaka University, Suita 565-0871, Japan

