## Supplementary Figures for "Quantification of the escape from X chromosome inactivation with the million cell-scale human single-cell omics datasets reveals heterogeneity of escape across cell types and tissues"

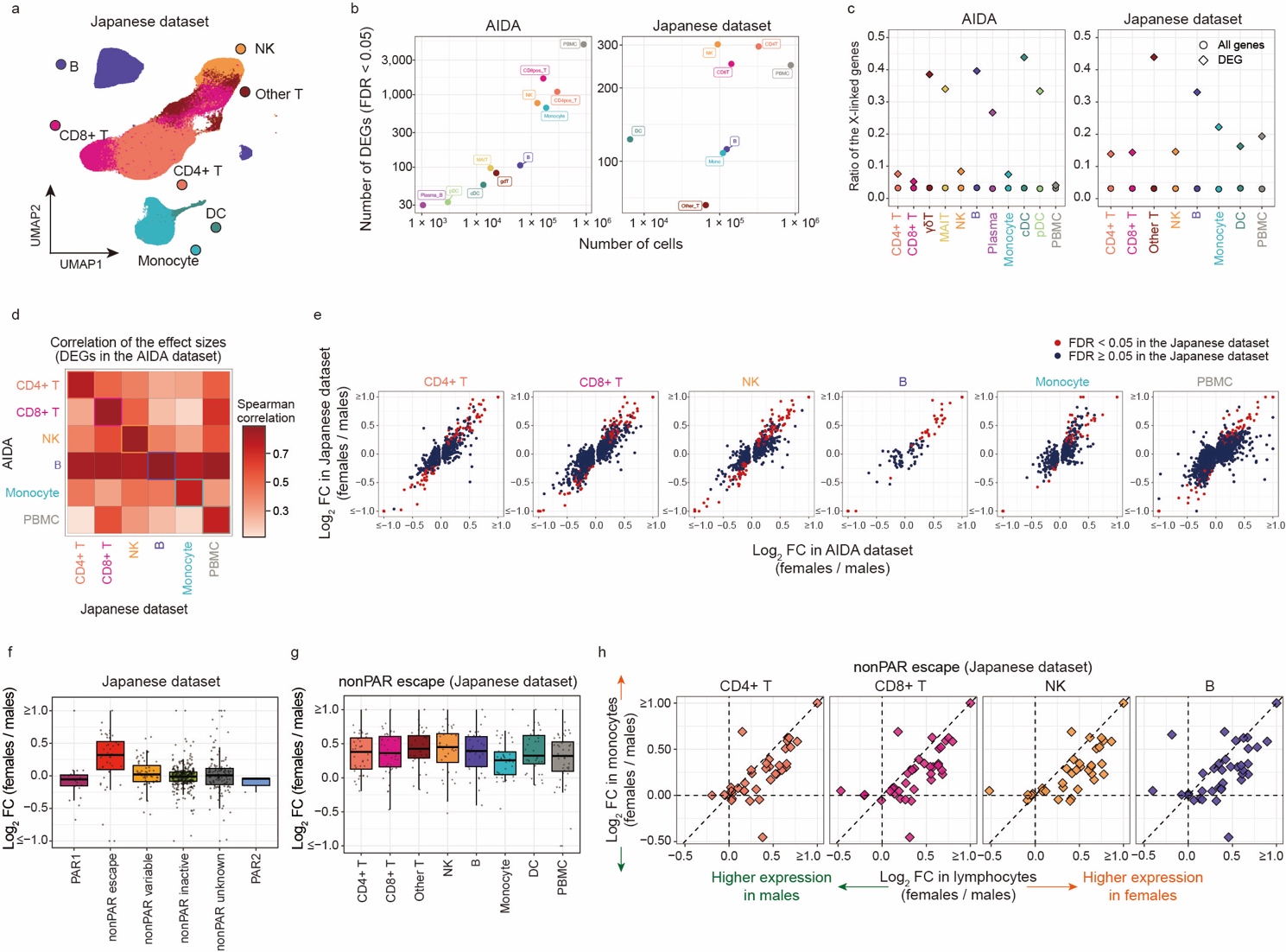

**Supplementary Figure 1. The results of the pseudobulk differentially expressed gene analysis are consistent between the AIDA and Japanese datasets**

**a,** UMAP of the Japanese dataset. **b,** The relationship between the number of cells (x-axis) and the number of the significant DEGs (FDR < 0.05; y-axis) for the AIDA (left) and Japanese (right) datasets. The colors indicate the cell types. **c,** The ratios of the X-linked genes among all genes (circle) and DEGs (rhombus) are indicated for the AIDA (left) and Japanese (right) datasets. The colors indicate the cell types. **d,** Spearman correlations between the effect sizes in the DEG analysis for the AIDA (y-axis) and Japanese (x-axis) datasets for DEGs (AIDA dataset) in the major cell types. **e,** Scatter plots represent the effect sizes of the significant DEGs detected in the AIDA dataset in the DEG analysis for the AIDA (x-axis) and Japanese (y-axis) datasets. The colors of the points represent whether the genes are significant DEGs in the Japanese dataset. **f,** A box plot represents log2 fold-changes of the gene expression between sexes in the Japanese dataset. Genes are grouped according to the XCI status annotated in the previous study. The boxplot indicates the median values (center lines) and IQRs (box edges), with the whiskers extending to the most extreme points within the range between (lower quantile − [1.5  ×  IQR]) and (upper quantile  +  [1.5  ×  IQR]). **g,** A box plot represents log2 fold-changes of the escapee gene expression between sexes across cell types in the Japanese dataset. The boxplot indicates the median values (center lines) and IQRs (box edges), with the whiskers extending to the most extreme points within the range between (lower quantile − [1.5  ×  IQR]) and (upper quantile  +  [1.5  ×  IQR]). **h,** Scatter plots represent pairwise comparisons of the log2 fold-changes of the escapee gene expression between sexes in the Japanese dataset. The y-axes represent the log2 fold-changes in monocytes and the x-axes represent the log2 fold-changes in lymphocytes. The dashed lines represent *x* = 0, *x* = *y*, and *y* = 0. DEG, differentially expressed genes; AIDA, Asian Immune Diversity Atlas; FC, fold-changes; FDR, false discovery ratio; IQR, interquartile range; PAR, pseudoautosomal region; PBMC, peripheral blood mononuclear cells; scRNA-seq, single-cell RNA-seq; UMAP, Uniform manifold approximation and projection; XCI, X chromosome inactivation.

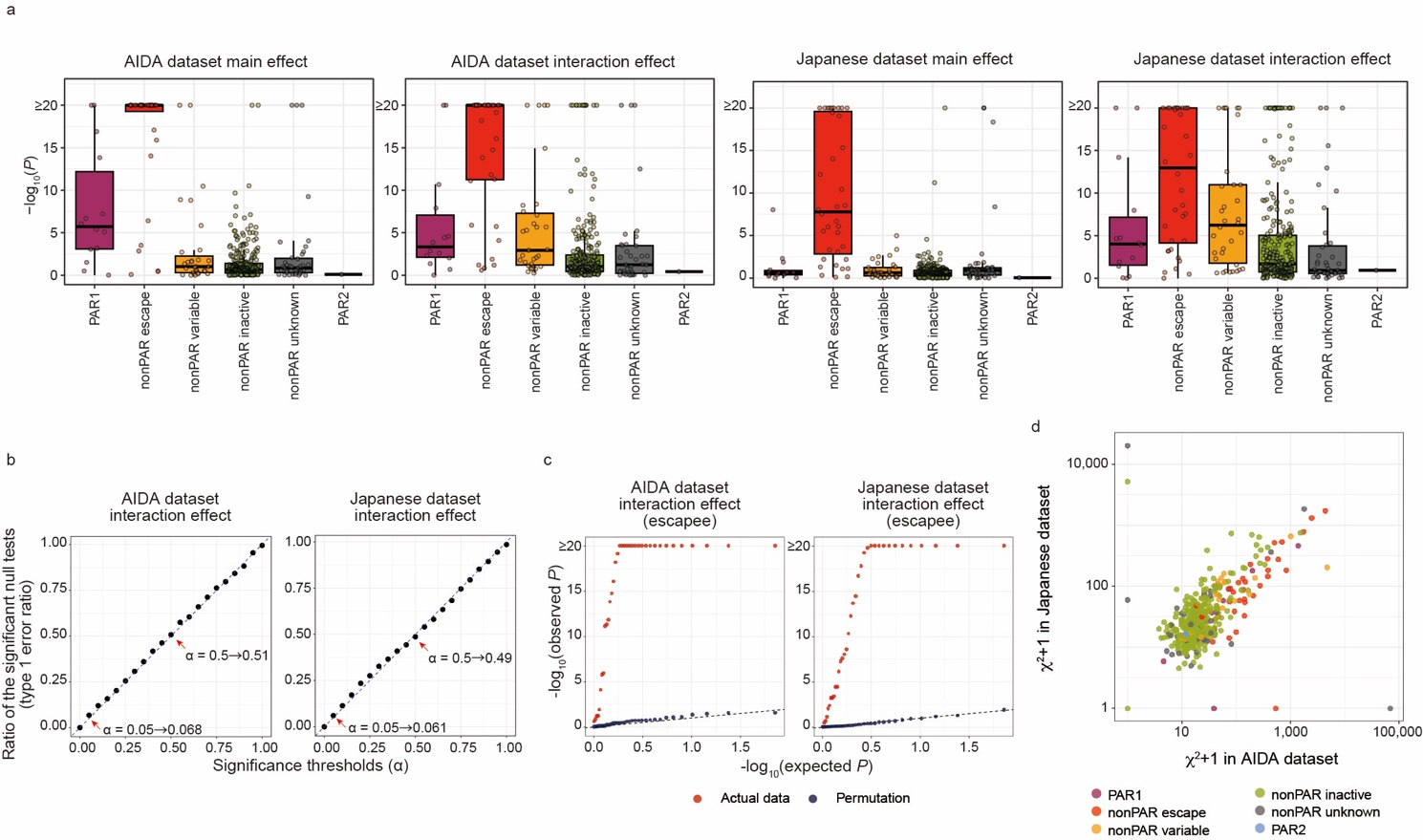

**Supplementary Figure 2. The results of the single-cell level differentially expressed gene analysis are consistent between the AIDA and Japanese datasets**

**a,** A box plot represents P-values for the sex term (AIDA dataset), sex × cell state term (AIDA dataset), sex term (Japanese dataset), and sex × cell state (Japanese dataset) in the single-cell level DEG analysis. Genes are grouped according to the XCI status annotated in the previous study. The boxplot indicates the median values (center lines) and IQRs (box edges), with the whiskers extending to the most extreme points within the range between (lower quantile − [1.5  ×  IQR]) and (upper quantile  +  [1.5  ×  IQR]). **b,** Ratio of significant tests under cell state permutation in the AIDA and Japanese datasets. Each dot represents the ratio of the significant tests (y-axis) at the given alpha threshold (x-axis). **c,** Q-Q plots for the P-values for the sex × cell state term of escapee genes (left, AIDA dataset; right, Japanese dataset). The color of the dots represents whether the test is performed for the actual data or under cell state permutation. **d,** A scatter plot represents the relationship between the χ2 statistics for the sex × cell state term in the AIDA (x-axis) and Japanese dataset (y-axis). The colors of the points represent the XCI status annotated in the previous study. AIDA, Asian Immune Diversity Atlas; DEG, differentially expressed genes; IQR, interquartile range; PAR, pseudoautosomal region; scRNA-seq, single-cell RNA-seq; XCI, X chromosome inactivation.
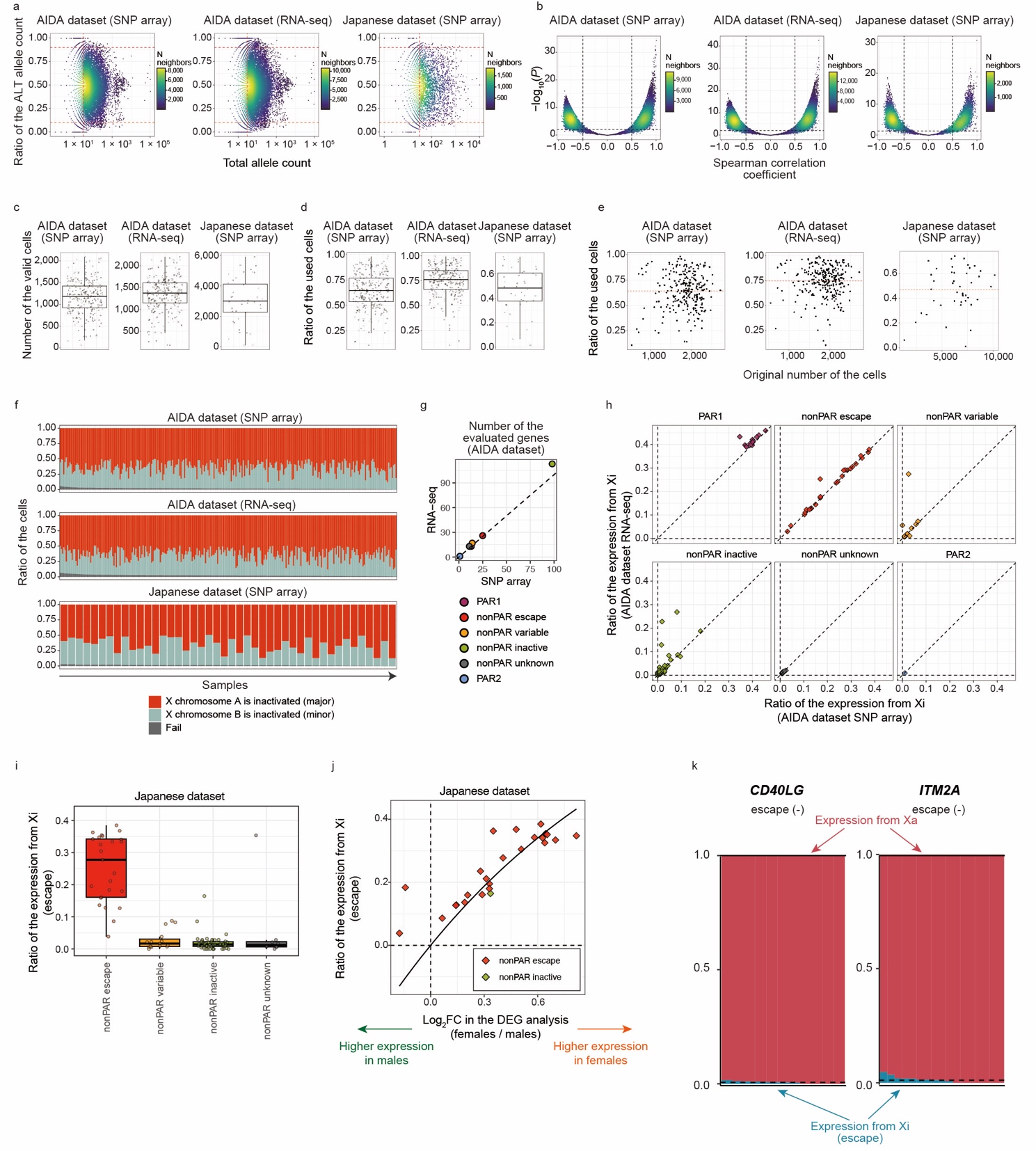

**Supplementary Figure 3. Quantification of the escape from XCI by scLinaX**

**a,** The relationships between the ratio of the ALT allele counts (y-axis) and the total allele counts (x-axis) for each SNP for each dataset. The points represent the pairs of sample and SNP, and the colors of the points represent the density of the neighboring points. The red dashed lines indicate the thresholds for QC. **b,** The relationships between the -log_10_ (P-value) (y-axis) and correlation coefficients (x-axis) of the Spearman correlation tests for the pseudobulk ASE profiles generated during scLinaX workflow. The points represent the pairs of two pseudobulk profiles, and the colors of the points represent the density of the neighboring points. The dashed lines indicate the thresholds for QC. **c,d,** The boxplots represent the number of valid cells (c) and the ratio of the used cells (cells for which any of the reference SNPs are detected; d) for each sample. The boxplot indicates the median values (center lines) and IQRs (box edges), with the whiskers extending to the most extreme points within the range between (lower quantile − [1.5  ×  IQR]) and (upper quantile  +  [1.5  ×  IQR]). **e,** The relationship between the original number of the cells (x-axis) and the ratio of the used cells (y-axis). The points represent samples. The red dashed lines indicate the mean ratio of the used cells across samples. **f,** A bar plot represents the ratio of the cells that different X chromosomes are inactivated or are removed from the analysis due to the bi-allelic expression of the reference SNPs. **g,** Number of the genes evaluated in the scLinaX analysis of the AIDA dataset with the SNP data based on the SNP array (x-axis) and called from scRNA-seq data (y-axis). The colors of the dots represent the XCI status annotated in the previous study. **h,** Plots represent the ratio of the expression from Xi in the AIDA dataset calculated from the SNP data derived from SNP array data (x-axis) and scRNA-seq data (y-axis). Genes are grouped according to the XCI status annotated in the previous study. **i,** A box plot represents the estimated ratio of the expression from Xi in the Japanese dataset. Genes are grouped according to the XCI status annotated in the previous study. The boxplot indicates the median values (center lines) and IQRs (box edges), with the whiskers extending to the most extreme points within the range between (lower quantile − [1.5  ×  IQR]) and (upper quantile  +  [1.5  ×  IQR]). **j,** A plot represents the relationship between the log2 fold-changes in the DEG analysis (x-axis) and the ratio of the expression from Xi (y-axis) in the Japanese dataset. Genes that are annotated as escape genes or shows evidence of escape in the scLinaX analysis (*SEPTIN6*) are indicated. The curved line indicates the theoretical relationship under the assumption that differential gene expression between sexes is solely due to the expression from Xi and total gene expression in males and Xa-derived gene expression in females are at the same level. **k,** Plots represent the ratio of the expression from Xa and Xi at an individual level for the *CD40LG* and *ITM2A* gene. The dashed horizontal line represents the mean ratio of the expression from Xi across samples. Since SNPs on the *ITM2A* gene were included in the initial analysis of the Japanese dataset, scLinaX analysis removing reference SNPs on the *ITM2A* genes was specifically performed for making the plot (right). AIDA, Asian Immune Diversity Atlas; ALT, alternative allele; ASE, allele-specific expression; CI, confidence interval; DEG, differentially expressed genes; FC, fold-changes; FDR, false discovery ratio; IQR, interquartile range; PAR, pseudoautosomal region; QC, quality control; REF, reference allele; SNP, single nucleotide polymorphism; Xa, active X chromosome; XCI, X chromosome inactivation; Xi, inactive X chromosome.

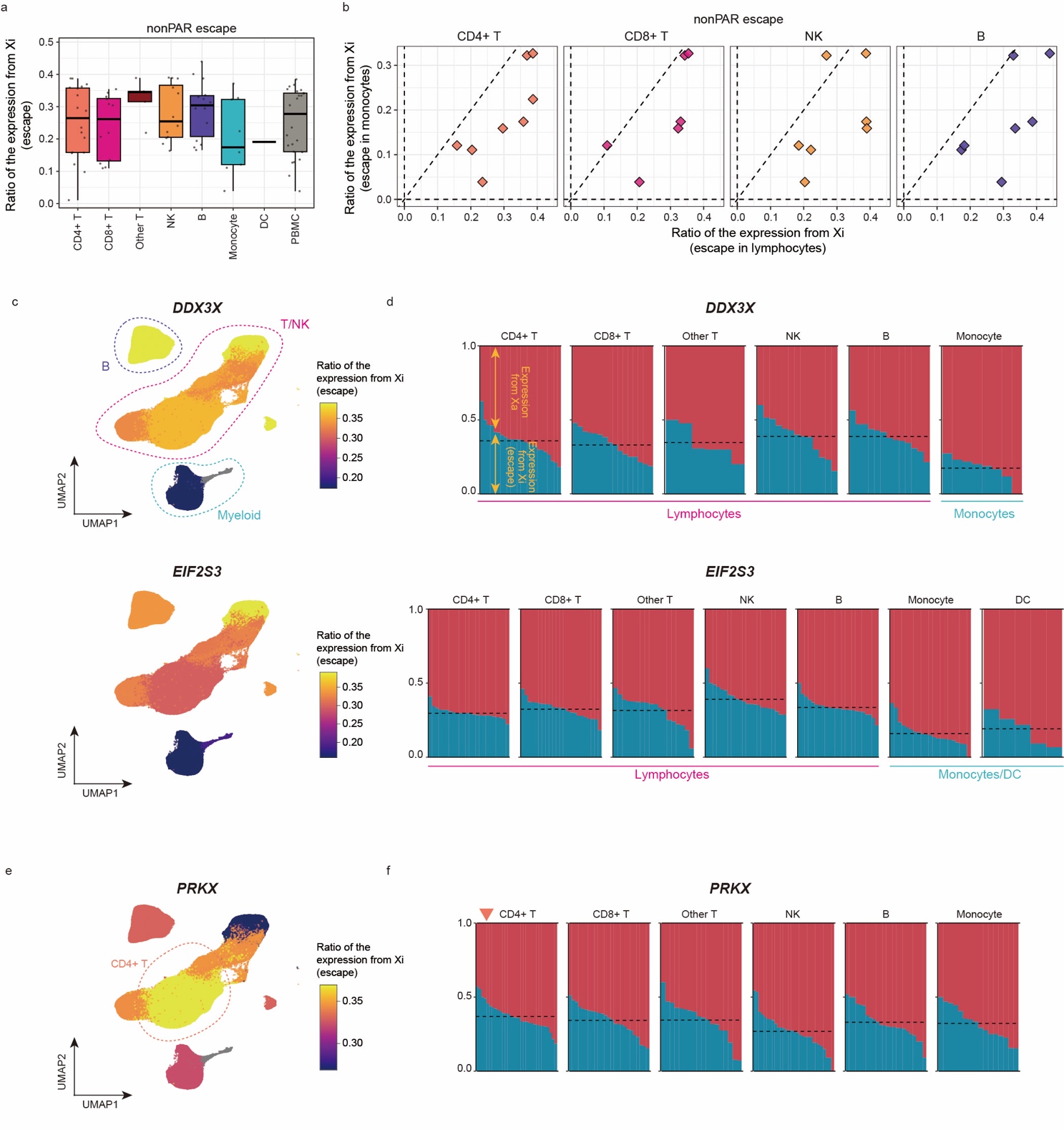

**Supplementary Figure 4. The scLinaX-based quantification of the escape from XCI across immune cell types in the Japanese dataset**

**a,** A box plot represents the estimated ratio of the expression from Xi for escapee genes across cell types in the Japanese dataset. The boxplot indicates the median values (center lines) and IQRs (box edges), with the whiskers extending to the most extreme points within the range between (lower quantile − [1.5  ×  IQR]) and (upper quantile  +  [1.5  ×  IQR]). **b,** Scatter plots represent pairwise comparisons of the ratio of the expression from Xi for escapee genes in the Japanese dataset. The y-axes represent the ratio of the expression from Xi in monocytes and the x-axes represent the ratio of the expression from Xi in lymphocytes. The dashed lines represent *x* = 0, *x* = *y*, and *y* = 0. **c,** UMAPs of the Japanese dataset colored according to the ratio of the expression from Xi estimated for each cell type. Examples of genes that show a higher ratio of expression from Xi in lymphocytes than monocytes are indicated. Cell types whose ratio of the expression from Xi could not be estimated are colored grey. **d,** Plot represents the ratio of the expression from Xa and Xi at an individual level for each cell type in the Japanese dataset. Examples of genes that show a higher ratio of expression from Xi in lymphocytes than monocytes, *DDX3X* and *EIF2S3* genes, are indicated. The dashed horizontal line represents the mean ratio of the expression from Xi across samples for each cell type. **e,** A UMAP of the Japanese dataset colored according to the ratio of the expression from Xi estimated for each cell type. The *PRKX* gene, which shows a unique pattern of heterogeneity of the escape across cell types, is indicated. Cell types whose ratio of the expression from Xi could not be estimated are colored grey. **f,** Plot represents the ratio of the expression from Xa and Xi at an individual level for each cell in the Japanese dataset. The *PRKX* gene, which shows a unique pattern of heterogeneity of the escape across cell types, is indicated. The dashed horizontal line represents the mean ratio of the expression from Xi across samples for each cell type. IQR, interquartile range; UMAP, Uniform manifold approximation and projection; Xa, active X chromosome; XCI, X chromosome inactivation; Xi, inactive X chromosome.

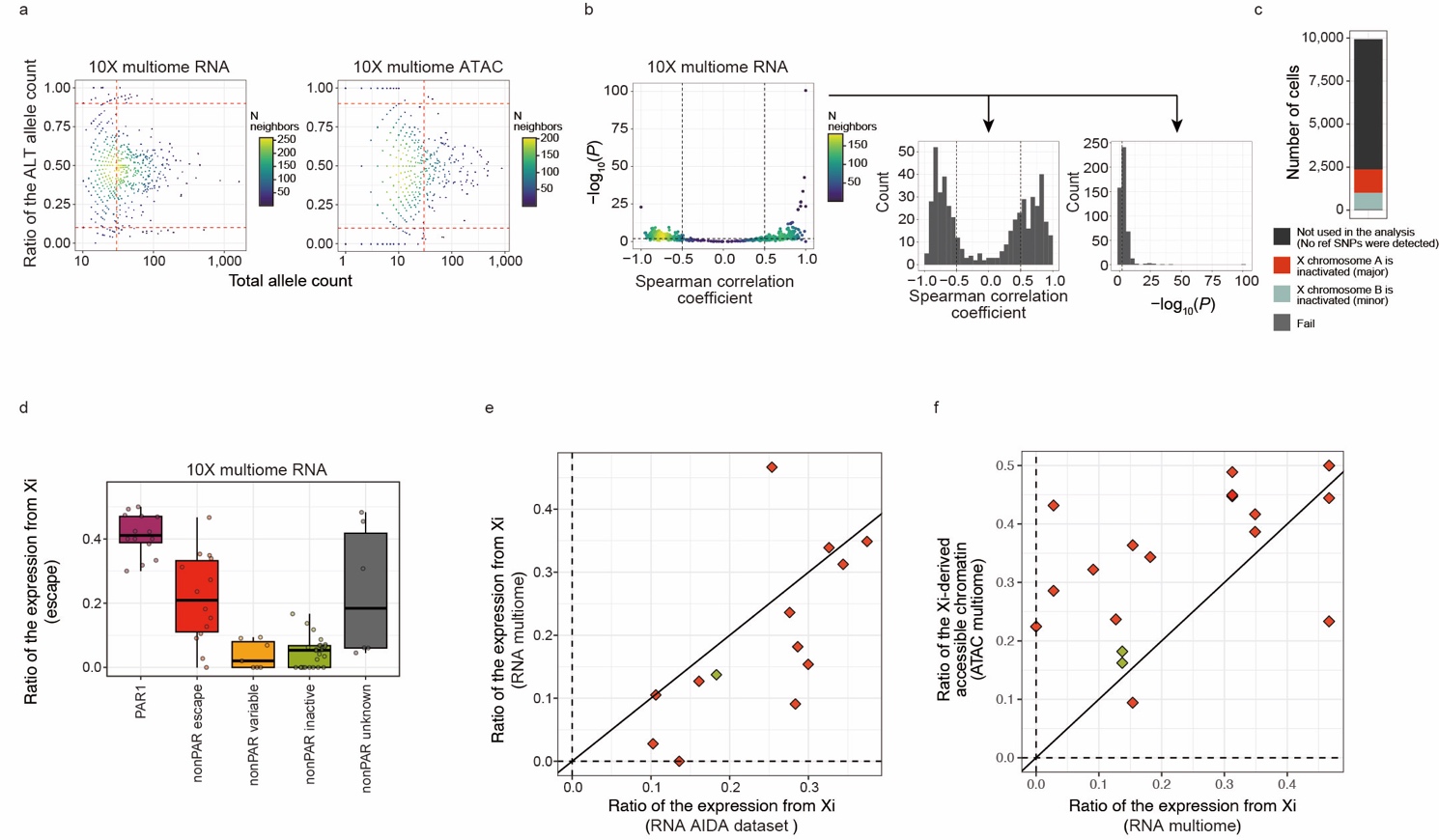

**Supplementary Figure 5. Application of scLinaX-multi to the 10X multiome dataset**

**a,** The relationships between the ratio of the ALT allele counts (y-axis) and the total allele counts (x-axis) for each SNP for each modality. The points represent the pairs of sample and SNP, and the colors of the points represent the density of the neighboring points. The red dashed lines indicate the thresholds for QC. **b,** The relationships between the -log_10_ (P-value) (y-axis) and correlation coefficients (x-axis) for the Spearman correlation tests for the pseudobulk ASE profiles generated during scLinaX-multi workflow (left). The points represent the pairs of two pseudobulk profiles, and the colors of the points represent the density of the neighboring points. The dashed lines indicate the thresholds for QC. Histograms for the -log_10_(P-value) (middle) and correlation coefficients (right) of the Spearman correlation tests are also indicated. **c,** A bar plot represents the number of the cells that are not used for the analysis (black), different X chromosomes are inactivated (red/blue), or are removed from the analysis due to the bi-allelic expression of the reference SNPs (grey). **d,** A box plot represents the estimated ratio of the expression from Xi for the gene expression data of the multiome dataset. Genes are grouped according to the XCI status annotated in the previous study. The boxplot indicates the median values (center lines) and IQRs (box edges), with the whiskers extending to the most extreme points within the range between (lower quantile − [1.5  ×  IQR]) and (upper quantile  +  [1.5  ×  IQR]). **e,** A plot represents the concordance of the ratio of the expression from Xi between the AIDA dataset (x-axis) and the multiome dataset (RNA; y-axis). Genes that are annotated as escape genes or shows evidence of escape in the scLinaX analysis (*SEPTIN6*) are indicated. The black line indicates *x* = *y*. **f,** A plot represents the relationship between the ratio of the expression from Xi (multiome, RNA-level, x-axis) and the ratio of the accessible chromatin derived from Xi (y-axis) for each peak–nearest gene pair. Genes that are annotated as escape genes or shows evidence of escape in the scLinaX analysis (*SEPTIN6*) are indicated. The black line indicates *x* = *y*. AIDA, Asian Immune Diversity Atlas; ALT, alternative allele; ASE, allele-specific expression; ATAC, Assay for Transposase-Accessible Chromatin; IQR, interquartile range; PAR, pseudoautosomal region; QC, quality control; REF, reference allele; SNP, single nucleotide polymorphism; Xa, active X chromosome; XCI, X chromosome inactivation; Xi, inactive X chromosome.

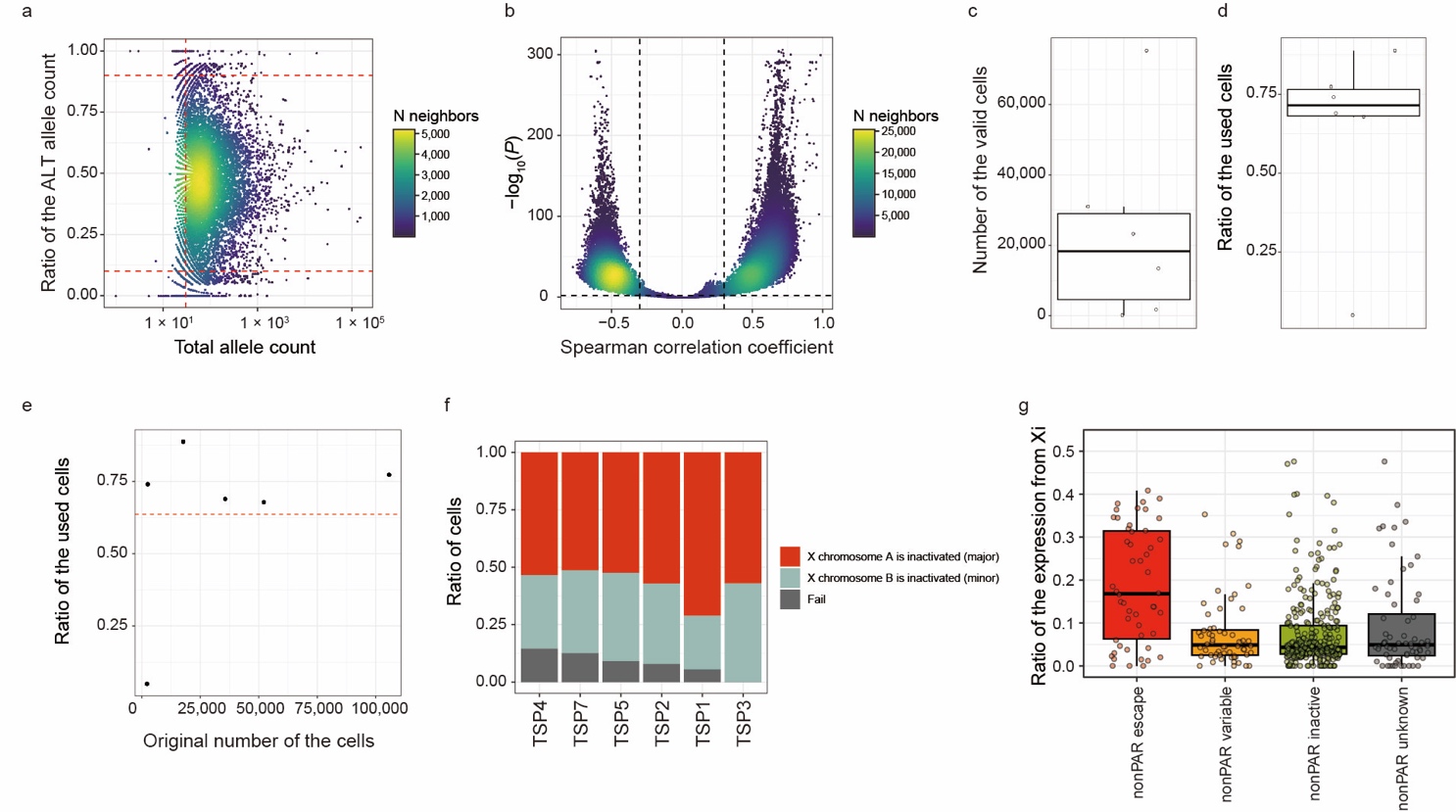

**Supplementary Figure 6. Application of scLinaX to the Tabula Sapiens dataset**

**a,** The relationships between the ratio of the ALT allele counts (y-axis) and the total allele counts (x-axis) for each SNP. The points represent the pairs of sample and SNP, and the colors of the points represent the density of the neighboring points. The red dashed lines indicate the thresholds for QC. **b,** The relationships between the -log_10_ (P-value) (y-axis) and correlation coefficients (x-axis) of the Spearman correlation tests for the pseudobulk ASE profiles generated during scLinaX workflow. The points represent the pairs of two pseudobulk profiles, and the colors of the points represent the density of the neighboring points. The dashed lines indicate the thresholds for QC. **c,d,** The boxplots represent the number of valid cells (c) and the ratio of the used cells (cells for which any of the reference SNPs are detected; d) for each sample. The boxplot indicates the median values (center lines) and IQRs (box edges), with the whiskers extending to the most extreme points within the range between (lower quantile − [1.5  ×  IQR]) and (upper quantile  +  [1.5  ×  IQR]). **e,** The relationship between the original number of the cells (x-axis) and the ratio of the used cells (y-axis). The points represent samples. The red dashed lines indicate the mean ratio of the used cells across samples. **f,** A bar plot represents the ratio of the cells that different X chromosomes are inactivated or are removed from the analysis due to the bi-allelic expression of the reference SNPs. **g,** A box plot represents the estimated ratio of the expression from Xi. Genes are grouped according to the XCI status annotated in the previous study. The boxplot indicates the median values (center lines) and IQRs (box edges), with the whiskers extending to the most extreme points within the range between (lower quantile − [1.5  ×  IQR]) and (upper quantile  +  [1.5  ×  IQR]). ALT, alternative allele; ASE, allele-specific expression; IQR, interquartile range; PAR, pseudoautosomal region; QC, quality control; REF, reference allele; SNP, single nucleotide polymorphism; Xa, active X chromosome; XCI, X chromosome inactivation; Xi, inactive X chromosome.

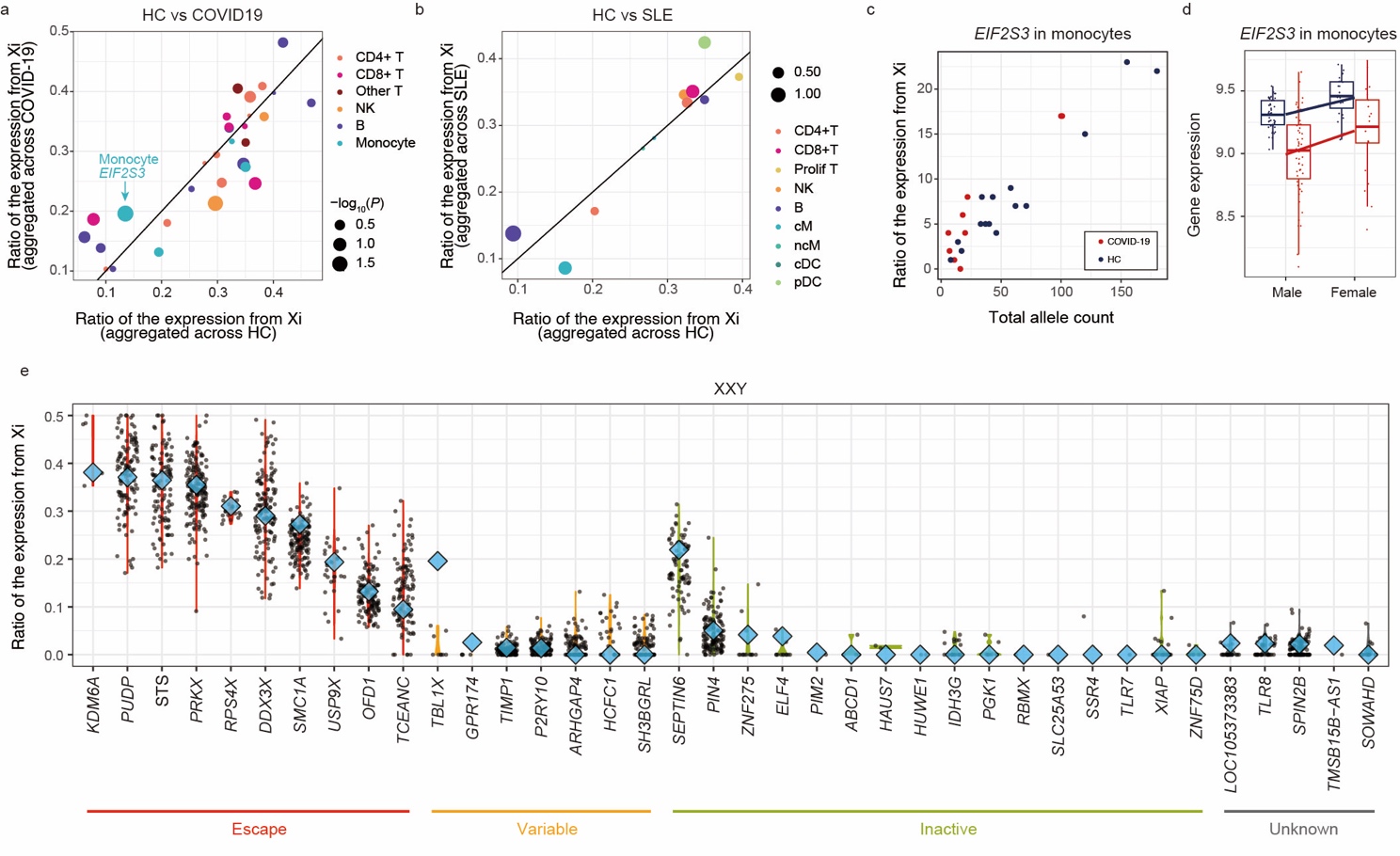

**Supplementary Figure 7. Evaluation of the escape in disease conditions**

**a,b,** The ratio of the expression from Xi in healthy subjects (x-axis) and disease patients (y-axis; a, COVID-19; b, SLE). Each point represents a pair of genes and cell types. The colors and sizes of the points indicate the cell type and P-values. The line represents *x* = *y*. **c,** The relationship between the ratio of the expression from Xi (y-axis) and total allele count (x-axis) for the *EIF2S3* gene in monocyte. Each dot represents each sample. The color of the points represents the disease affection status of the samples. **d,** A boxplot represents the expression of the *EIF2S3* gene in monocyte. The x-axis indicates the sex and the color of the plots represents the disease affection status. The boxplot indicates the median values (center lines) and IQRs (box edges), with the whiskers extending to the most extreme points within the range between (lower quantile − [1.5  ×  IQR]) and (upper quantile  +  [1.5  ×  IQR]). **e,** The ratio of the expression from Xi in a male sample with a karyotype of XXY is indicated as blue rhombuses. A violin plot represents the ratio of the expression from Xi in the AIDA datasets. AIDA, Asian Immune Diversity Atlas; COVID-19, coronavirus disease of 2019; HC, healthy control; IQR, interquartile range; PAR, pseudoautosomal region; REF, reference allele; SLE, systemic lupus erythematosus; SNP, single nucleotide polymorphism; Xa, active X chromosome; XCI, X chromosome inactivation; Xi, inactive X chromosome.

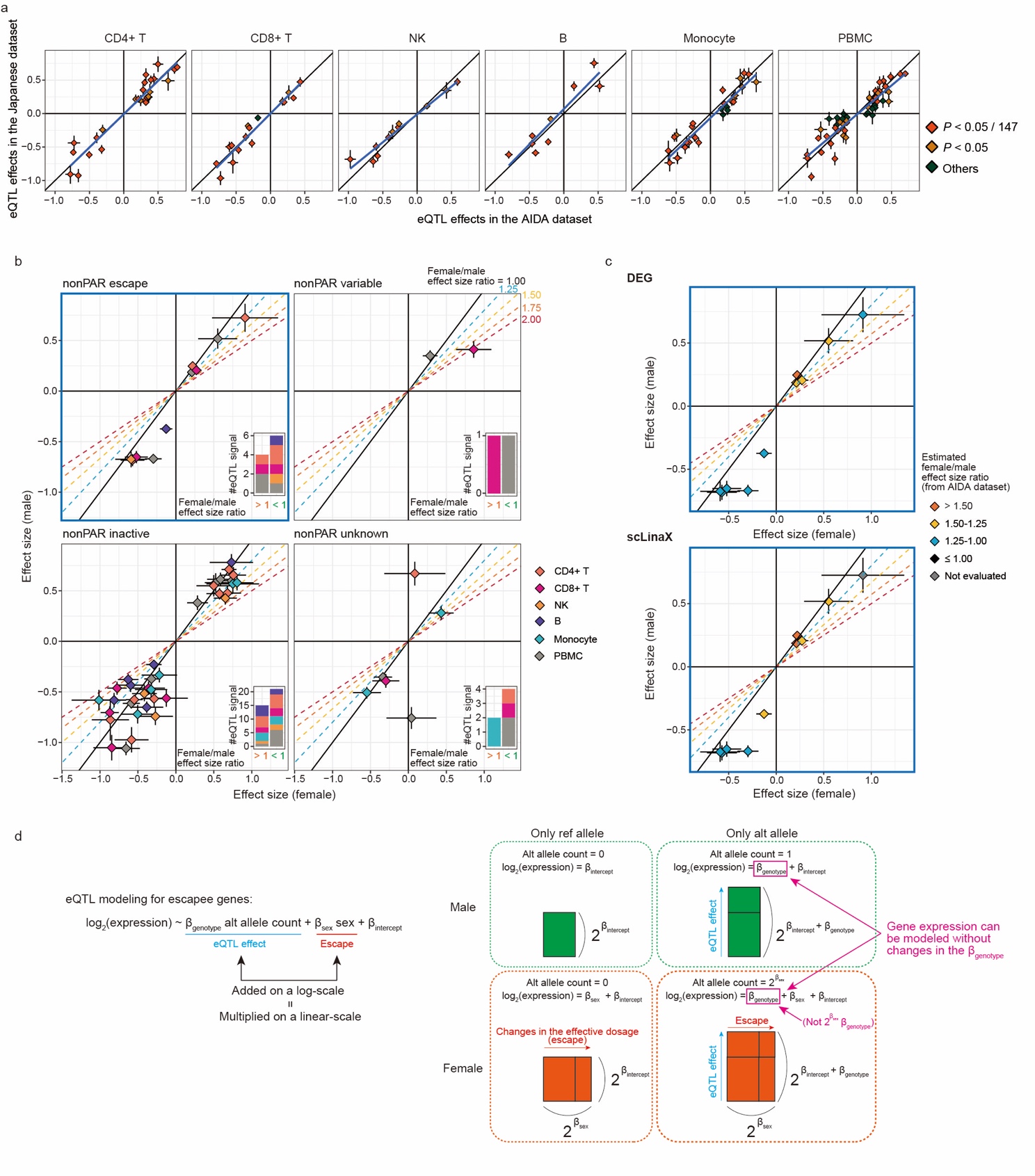

**Supplementary Figure 8. Comparison of the eQTL effect sizes between sexes**

**a,** The relationship between the eQTL effect sizes between the AIDA dataset (x-axis) and Japanese dataset (y-axis) for the significant eQTL signals detected in the AIDA dataset. The color of the points represents the significance of the eQTL effects in the Japanese dataset (P < 5 × 10^-8^). The blue line indicates the regression line. **b,** Scatter plots represent the effect sizes of the significant eQTL signals (P < 5 × 10^-8^; AIDA dataset) in the female-only (x-axis) and male-only (y-axis) analyses with the Japanese dataset, separately for each XCI status. The error bars indicate standard errors. The color of the plots indicates the cell type in which the eQTL signals are identified. The oblique lines correspond to the female/male effect size ratios described in the plots. The attached bar plots indicate the number of eQTL signals that have larger effect sizes in females (left) and males (right). **c,** The scatter plots for escapee genes (b, upper left) were colored according to the estimated female/male effect size ratio based on the DEG analysis (top) and scLinaX analysis (bottom) with the AIDA dataset. Genes that are not evaluated in the scLinaX analyses are colored grey. **d,** A schematic illustration of the effect of the gene expression normalization method on the eQTL analysis of the escapee genes. DEG, differentially expressed genes; eQTL, expression quantitative trait locus; PAR, pseudoautosomal region; XCI, X chromosome inactivation.
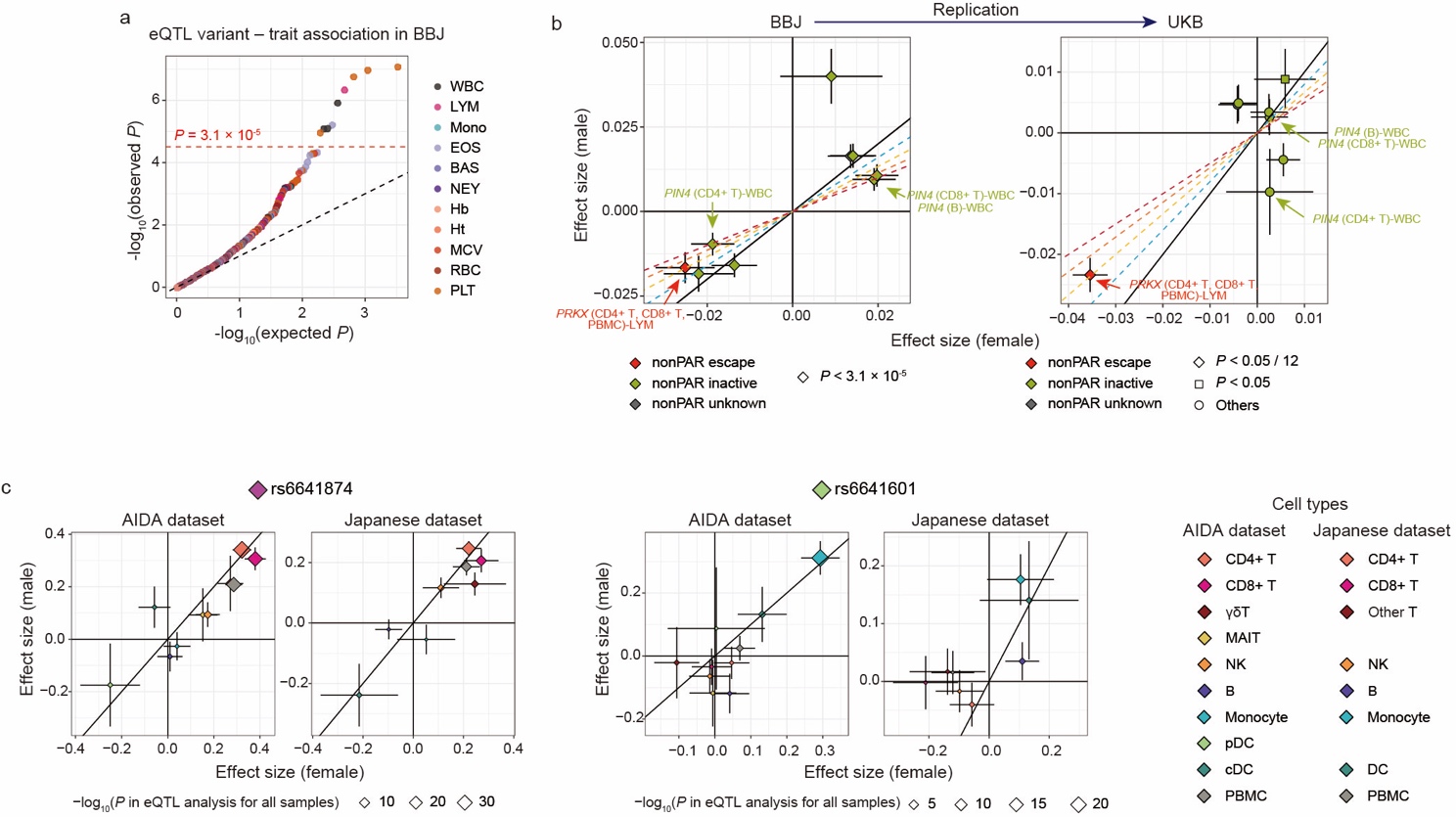

**Supplementary Figure 9. Comparison of the blood-related trait QTL analysis effect sizes between sexes**

**a,** A Q-Q plot represents the association between the significant eQTL variants detected in the AIDA dataset analysis and the blood-related traits in the BBJ cohort. The color of the dots represents the blood-related phenotypes. The red dashed line represents the significance threshold under a multiple-test correction. **b,** The comparisons of the blood-related trait QTL analysis effect sizes between sexes in the BBJ (left) and UKB (right) cohort. The variant–phenotype association swhich satisfy the significance threshold in (a) are indicated. The color of the plots represents the XCI status annotated in the previous study. The shapes of the points in the right panel (UKB) indicate whether the significant variant–phenotype association in BBJ is replicated with the UKB dataset. The error bars indicate standard errors. **c,** Scatter plots represent the effect sizes of the two eQTL signals (rs6641874–*PRKX* and rs6641601–*PRKX*) in the female-only (x-axis) and male-only (y-axis) analyses with the AIDA and Japanese dataset. The error bars indicate standard errors. The color of the points indicates the cell type in which the eQTL signals are identified. The size of the points indicates the P-values in the eQTL analysis with all samples. BBJ, BioBank Japan; PAR, pseudoautosomal region; QTL, quantitative trait locus; UKB, UK Biobank.
